# Analysis of Biological Record Data: Improvements of the Occupancy–Detection Models

**DOI:** 10.1101/408724

**Authors:** Shiyu Li

## Abstract

Many indicators are used to monitor the progress of the target which aims to stop the biodiversity loss by 2020. However, the occupancy-detection model which is currently applied to calculate the indicator is biased. Hence, more robust models are required to track the trend of the species precisely. This research first reviews the previous works on improving this occupancy-detection model by changing the prior distributions of one of the quantities and of the models considered previously, a model based on a random walk is found to be the most appropriate although it has some potential deficiencies. Then this research provides some potential improvements of the random walk model by changing the way of modelling the prior distributions of each quantity and changing the model structure. Then the hoverflies datasets are used in this research to analyse the performance of the models. These models are compared by the running times of fitting the models and the plots of the trend of the species of all models. As a result, the categorical list length model is considered to be the most precise model among all models with a reasonable running time. Then, we fit this model with a large dataset, however, it takes a long running time to get the result. Finally, some potential improvements are suggested which may be useful for further research.

## 1 Introduction

The aim of this research is to find improved methods for estimating changes over time in biodiversity as represented by species’ ranges, using records provided by volunteers. This chapter will introduce the background and motivation for the research. Then we will provide information of the species studied in this research and describe the dataset used. Finally, there will be an overview of this research.

### 1.1 Background

The Convention on Biological Diversity which is a treaty proposed by the United Nations has set twenty targets to stop biodiversity loss by 2020 (Convention on Biological Diversity, 2010). These targets include the awareness of biodiversity and reasons causing biodiversity loss, halting biodiversity loss, sustainable production and consumption, ecosystem resilience, etc. To monitor the progress of the target, many indicators have been developed to track the trend of biodiversity. However, most of the indicators can only be applied to abundance dataset which are collected by national monitoring schemes. The abundance datasets are collected in a consistent and robust manner which are long term and have good geographical converge, and using statistical approaches to correct the biases (Burns et al., 2017). These data are ideal to be used in biodiversity indicators. However, these datasets are limited in a small number of species. Occurrence data is another type of data to be used to monitor the biodiversity. These data are collected by volunteers when they detect a species. They record this detection following a specific protocol (Outhwaite et al., 2017). However, the main problem of occurrence data is that substantial bias may be introduced in trend estimates because of the variation in recording activities (Isaac et al., 2014). Hence, Isaac et al. (2014) tests eleven models to find out the most robust and powerful model to analyse occurrence data. More information about this model will be provided in Chapter 2. However, this model is still not suitable for low recording intensity data of uncommon species. Therefore, Outhwaite et al. (2018) derives a model fitting the low recording intensity species. This process will be shown in Section 2.3.

### 1.2 Description of data

In this research, we use the hoverfly occurrence dataset in the UK which was provided by the British Hoverfly Recording Scheme at http://www.hoverfly.org.uk/ to evaluate the performance of each model. Hoverfly, also called flower fly, was named since they usually hover around flowers. Hoverfly is similar to bee with yellow markings but usually they do not sting or bite (Encyclopædia Britannica, 2018). It is more likely to find a hoverfly in farmlands and nature reserves than in urban areas (Baldock et al., 2015). The species we used here are anasimyia contracta, anasimyia interpuncta, episyrphus balteatus and eristalis pertinax respectively.

Anasimyia contracta prefers living in wetlands and ponds and its flight period is in summer (Speight, 2011). The distribution of this species is shown in Figure 1. Therefore, anasimyia contracta is a relatively common species in the UK, especially in the south part. Then, anasimyia interpuncta is a scarce species confined to the southeast of England. It often occurs associated with wetlands. Its flight period is also in summer time (Speight, 2011). The map of this species in the UK is shown in Figure 1 as well. Episyrphus balteatus is a relatively small hoverfly and almost ubiquitous (Speight, 2011). From Figure 1, episyrphus balteatus is widespread in the UK and it can be found almost everywhere, not only near to water. Finally, the eristalis pertinax, which can be found either in the wetland or in the forest, flies around bushes and settles on the ground by water. It is also a widespread species in the UK as shown in Figure 1.

**Figure 1.**
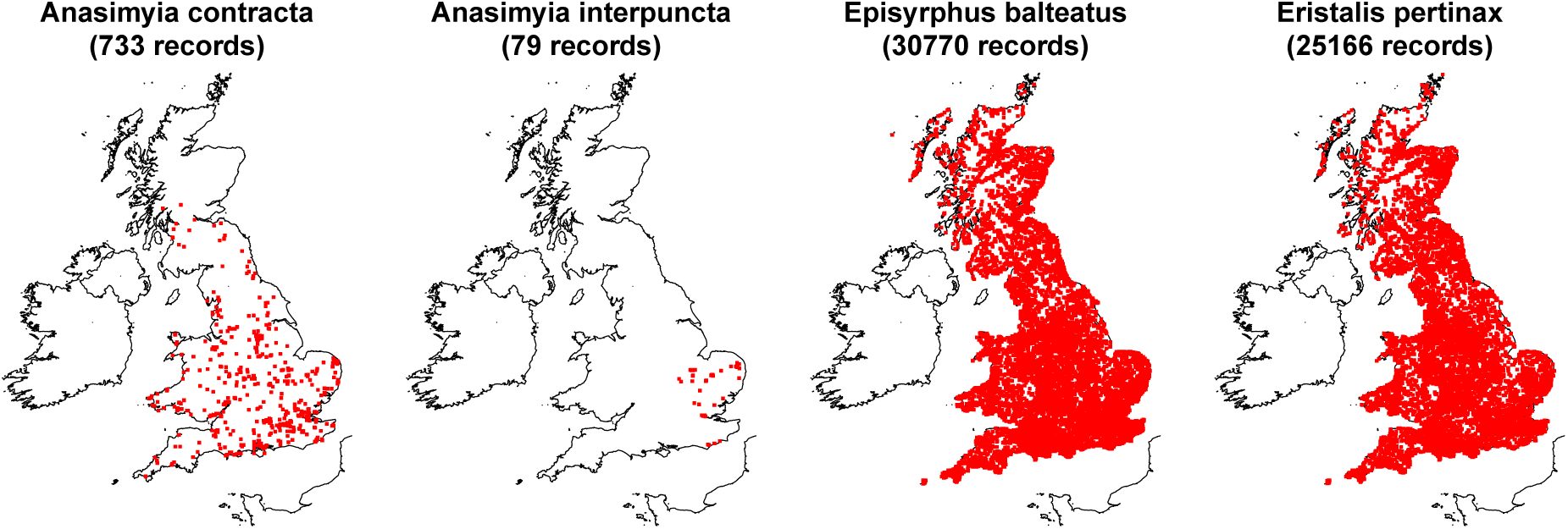
Distribution maps in the UK for four species from 1970-2014 (There is no data in Northern Ireland). Species names and number of records are shown on the top of each plot.

To test the performance of the model with low recording intensity data in this research, the data with anasimyia contracta and interpuncta will be applied in Chapter 5 since they have fewer records than the other two species. Also, the running time for each model will be less with fewer records. Then the data of episyrphus balteatus and eristalis pertinax will be applied in Chapter 6 using the best model to monitor the trends of these two species.

Each line in the dataset represents a single visit to a single location at a time. The location is recorded according to the site on a 1 *km*^2^ grid and the time is recorded with day, month and year. Each line in the dataset also contains the list length which means how many species was observed in each site and each day. Here we use list length to represent the effort of searching the species which is a common way to measure the effort (Szabo et al., 2010; Isaac et al., 2014). In the absence of explicit information on sampling effort, a longer list length can be taken as evidence that more efforts have been put into searching for species. Also, every line contains an indicator for whether or not the focal species was observed on the visit. Here, the focal species means the species we are concentrating on (which are anasimyia contracta, anasimyia interpuncta, episyrphus balteatus and eristalis pertinax respectively).

To gain a preliminary impression of the trend of each species, the proportion of detections among all visits for each species is plotted in Figure 2. This proportion of detection is calculated using the number of detections for each species divided by the number of visits each year, so the trend of the proportion of detection is usually similar to the trend of number of detections. However, we cannot use this method to estimate the trend of the species. Because the proportion are fluctuated violently from year to year which are not suitable for analysing of the state of the species. Also, this percentage is highly influenced by the number of visits each year. For example, Figure 2 shows a jump in the proportion in 2014 for all four species. However, this may be caused by fewer total visits in 2014 since the density of the number of detections shown in the histogram for these species are all reduced in 2014.

**Figure 2.**
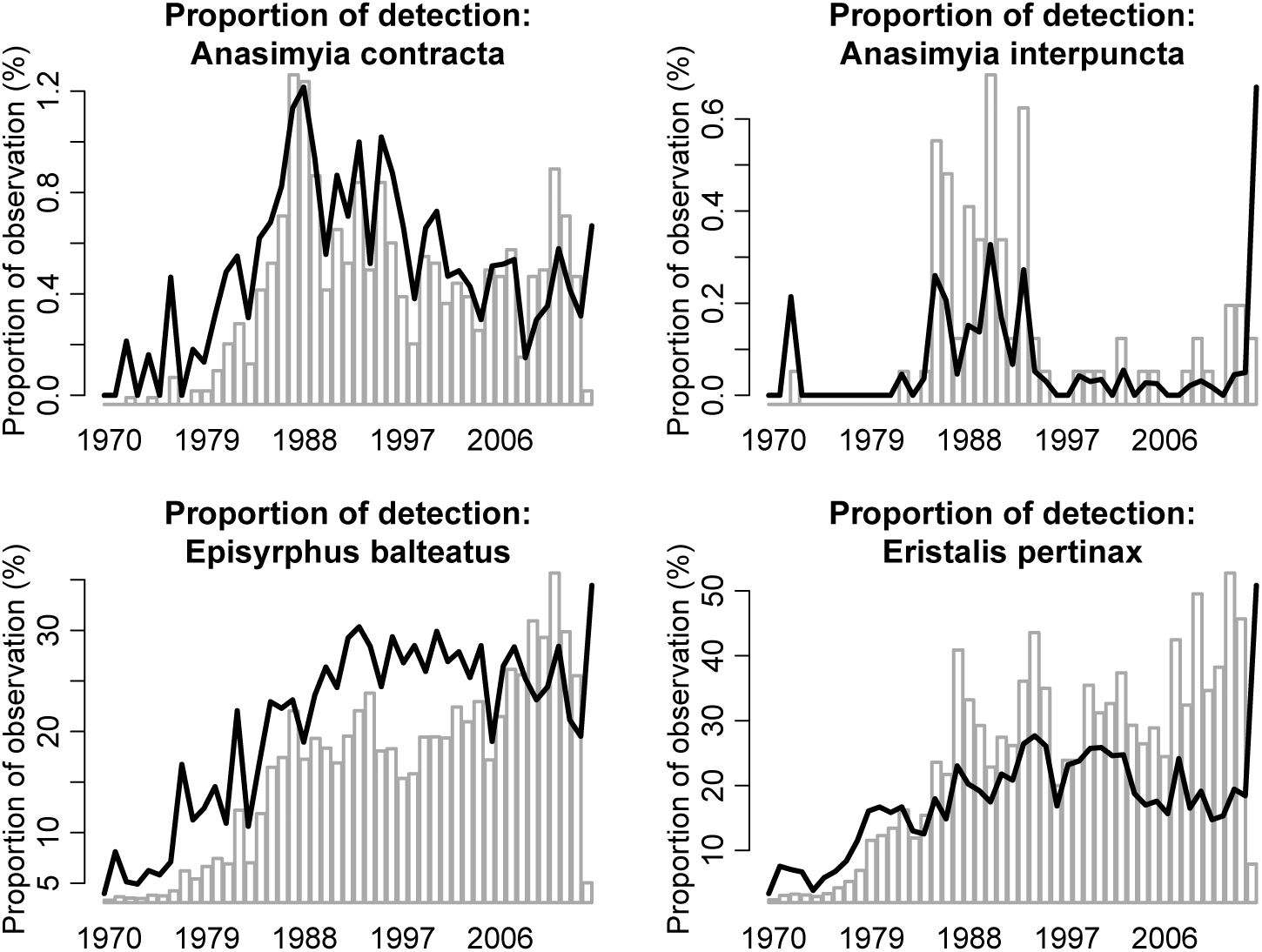
The name of each species are shown above each plot. The solid line in each plot shows, for each year, the number of visits for which the species in question was observed, expressed as a percentage of the total number of visits for that year. The histogram shows the density of the number of visits for which the species was observed each year.

However, we do not know the reasons of causing the decreasing in number of detections, whether it is because the decline of the species or less efforts in searching. Therefore, we need a more robust model to estimate the trend of a species. The model should be separated into two parts, one modelling the state of the species, i.e. whether it is present or not, the other modelling the detection of this species, i.e. if the species is present, whether it is reported or not.

### 1.3 Overview

This research aims to explore some alternative model formulations for analysing the kind of data shown in Figure 1. We first review the previous works in building these kinds of models, and we will subsequently consider to improve each part of this model and use the data of anasimyia contracta and anasimyia interpuncta to test the best model among all potential improvements. Finally, we will use this model to fit the species episyrphus balteatus and eristalis pertinax to see their trends in the past 20 years.

Chapter 2 provides a review of previous models that have been used to analyse data of this type. Chapter 3 then describes some potential improvements to the best model from Chapter 2. Chapter 4 contains the technical information of using Bayesian analysis to fit these models which are introduced in Chapter 2 and 3. Next, Chapter 5 compares these models from different aspects and finds the best model overall, and we will use this model in Chapter 6 to fit the data and get the trend of each species in the past 20 years. Finally, Chapter 7 provides conclusions about the findings in the research and potential improvements that would be beneficial to further investigations.

## 2 Occupancy-detection models

In this chapter we review the basic structure of occupancy-detection models for occurrence data, and present some of the specific models that have been suggested in the literature. As outlined in Section 1.2, for occurrence data, if there is no record of a species, we do not know whether it is because the species is not there or it is not recorded when it is present. To solve this problem, Isaac et al. (2014) uses the occupancy-detection model which is separated into two submodels, one occupancy model and one detection model.

The occupancy model describes the presence of the species, *z*_*it*_, of site i in year t. This is a binary variable, with 1 if present or 0 if absent. Let *Ψ*_*it*_ be the proportion that the site is occupied. Then the distribution of *z*_*it*_ is:

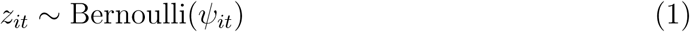

The detection model describes whether the species is recorded if it is present. Let *y*_*itv*_ be an detection indicator with 1 if recorded and 0 if not. Let *p*_*itv*_ denotes the probability that the species will be recorded on a single visit *v*, if the species is present at site *i* in year *t*. Then *y*_*itv*_ has a conditional distribution as follows:

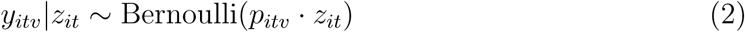

Here, the species can only be detected if it is present at site *i*. Hence, if *z*_*it*_ equals to zero, *y*_*itv*_ will also be zero. If the site is occupied however, the equation 2 gives*y*_*itv*_ *∼* Bernoulli(*p*_*itv*_).

Equations 1 and 2 form the basic structure of the occupancy-detection model. This chapter will outline the whole framework of the occupancy-detection model by Isaac et al. (2014) and the improvements by Outhwaite et al. (2018) together with the advantages or disadvantages of each model.

### 2.1 The base model

The base model we used here is from Outhwaite et al. (2018). As mentioned earlier in this chapter, the proportion of occupied sites and the probability of detection are used in the occupancy-detection model. Next, these probabilities are modelled in equations 3 and 4. The proportion of occupied sites *Ψ*_*it*_, representing the probability of the presence of the species, is modelled by a year effect *b*_*t*_ and a site effect *u*_*i*_ as following:

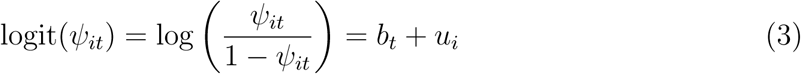

Then, the detection probability *p*_*itv*_ representing the probability of recording the species if present is modelled as following:

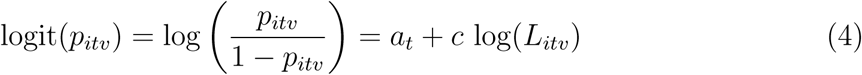

Here, *a*_*t*_ is a year effect, *L*_*itv*_ is the list length representing the total number of species recorded at visit *v* to site *i* in year *t*, and *c* is a parameter representing the relationship between the sampling effort and the detection probability of the focal species.

In equations 3 and 4, the only known quantity is the list length *L*_*itv*_. Since there are many unknown quantities, maximum likelihood estimation which is the most common approach to estimation and inference, is not suitable in this case. Therefore, Bayesian analysis is used here to fit the occupancy model.

The fundamental principle of Bayesian inference is to use the Bayes’ theorem to express the uncertainty about an unknown parameter vector *θ*, conditional on a vector of observed data *y*, via its posterior distribution *π*(*θ|y*) *∝π*(*y|θ*) *π*(*θ*), where *π*(*·*) denotes a generic probability density function. Here, the term *π*(*θ*) is the prior distribution for the unknown parameter, and represents the knowledge of the system in the absence of the data *y*. If we take an interval with required probability (e.g. 0.95) from the posterior distribution *π*(*θ|y*), this interval is called credible interval for *θ* which is the Bayesian ‘confidence’ interval.

The reason why Bayesian analysis works in this case is that modern Bayesian computational techniques can be used to carry out the inference in models with complex structures and many unknown parameters. However, standard techniques such as maximum likelihood are not feasible.

For Bayesian analysis, each unknown parameter must have a prior distribution describing the knowledge we know about the system. Therefore, the prior distribution of each parameter should be set before the data are imported. In the base model of Outh-waite et al. (2018), the priors were as follows, that most of the choices were taken from earlier work by Isaac et al. (2014).

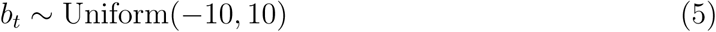

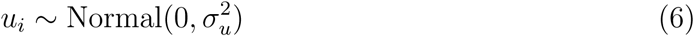

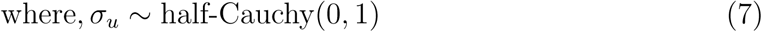

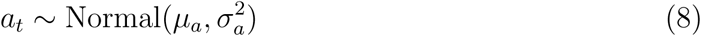

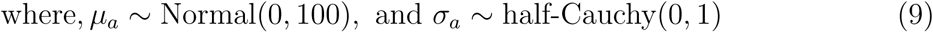

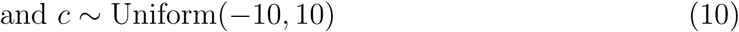

Here, the variances 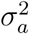 and 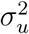 are specified via the inverses *τ*_*a*_ and *τ*_*u*_ which is computationally and mathematically convenient. The quantities 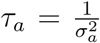and p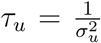are the precision parameters.

The priors of the year effect *a*_*t*_ and the site effect *u*_*i*_ have the hierarchical structures which mean the priors of these parameters also contain unknown parameters *µ*_*a*_, 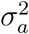 and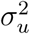, and these unknown parameters also have their own prior distributions (Gelman et al.,2013). The reason of using the hierarchical structure is that the hierarchical model is more flexible. For example, for the prior of the year effect *a*_*t*_, if the value of 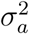is estimatedto be small, then the values of *a*_*t*_ will centre around the mean *µ*_*a*_. On the contrary, if the value of 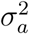 is estimated to be large, then the values of *a*_*t*_ generated will be almost the same as generating from the uniform distribution. Hence, the hierarchical structure allows the data to choose between these two situations by setting the prior of 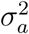 and this structure is therefore adapt to data.

However, unlike the parameters *a*_*t*_ and *u*_*i*_, the parameter *b*_*t*_ does not have a hierarchical prior structure with variance parameters *σ*^2^ assigned a prior distribution, which seems a bit inconsistent. Therefore, the parameter *b*_*t*_ does not have the flexibility to vary with the data. Also, if the data are not informative enough to overcome the prior, the precision of the estimates will be very low and the credible interval will be large. Outhwaite et al. (2018) does a simulation experiment to test whether this model is suitable for low recording intensity species. Based on the experiment, the base model has low precision with wide credible interval. Also, based on the bias level, the proportion of occupied sites estimated by this model can be highly under- or over-estimated. Therefore, the base model does not perform well for low recording intensity data.

### 2.2 Adaptive stationary model

As outlined in Section 2.1, the base model cannot adapt to the data. One improvement by Outhwaite et al. (2018) is changing the distribution of parameter *b*_*t*_ from uniform to normal distribution, with mean and variance as unknown parameters, which is similar to the year effect *a*_*t*_ in equations 8 and 9. Therefore, the equation 5 of *b*_*t*_ is replaced by the following:

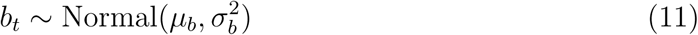

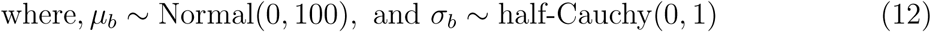

All other priors are not changed. Since the mean and variance of the parameter adapt to the data, the model is called adaptive stationary model.

In this model, as in the base model, the year effects *{b*_*t*_*}* are treated as independent from year to year *a priori*. However, in reality, if the species is present at a site during one year, it is also likely to be present in the following years. Also, based on the simulation experiment by Outhwaite et al. (2018), although adaptive stationary model has narrow credible interval, this model has great level of bias and tends to under-estimate the trend of the species. Therefore, the performance of adaptive stationary model is not well enough.

### 2.3 Random walk model

In the adaptive stationary model, the values of the year effects *b*_*t*_ are generated from the same prior distribution which may limit the rapid increase or decrease for the species. Alternatively, the values of the year effects *b*_*t*_ are modelled using different normal distributions in this section. As mentioned in Section 2.2, if a species is present during one year, it is likely to be present in the following year as well. Therefore, to make the year effect *b*_*t*_ depends on the previous year, this year effect is modelled in the form of *b*_*t*_ = *b*_*t−*1_ + E_*t*_, which is called random walk process. So this model is called random walk model. Then the equation 5 in the base model is replaced by the following:

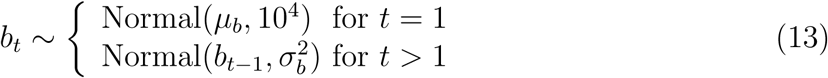

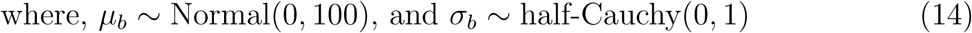

Here, the variance 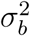 allows the year effect *b*_*t*_ to change based on the previous year. Large value of the variance means substantial fluctuations while the species fluctuated little will have small variance. By the simulation experiment in Outhwaite et al. (2018), the credible interval of random walk model is narrow. Also, this model has small bias. Therefore, random walk model performs well in this test.

### 2.4 Summary

This chapter reviews the occupancy-detection model constructed by Isaac et al. (2014) and improvements to fit the low recording intensity species by Outhwaite et al. (2018). Based on the problems of year effect *b*_*t*_ in the base model, the adaptive stationary model and the random walk model were constructed to improve it. A simulation experiment was done by Outhwaite et al. (2018) to test the performance of these models with low recording intensity data. The results of this simulation experiment suggested that among the three models considered, the random walk model delivers both the smallest bias and the narrowest credible intervals.

## 3 Further developments of the random walk model

The previous chapter reviewed some models for occupancy data, of which the random walk model was considered the most promising. However, there are opportunities to improve this model further: some of these opportunities are described in this chapter. As shown in equations 3 and 4, probabilities of occupancy and detection are modelled by year effects (*a*_*t*_ and *b*_*t*_), site effect (*u*_*i*_) and list length (*L*_*itv*_) with a parameter (*c*). Hence, in this chapter, we will consider potential improvements in each of these components of the model.

### 3.1 Improvements of year effect in occupancy submodel

We first consider the prior of year effect *b*_*t*_. In the prior of the random walk model, the mean of year effect *b*_*t*_ is equal to the previous one, *b*_*t−*1_, as shown in equation 13. However, there is another way of modelling the year effect *b*_*t*_ which is also considering its trend. More details about this model will be shown in section 3.1.1.

#### 3.1.1 Local linear trend model

In the random walk model, the year effect *b*_*t*_ is modelled by: *b*_*t*_ = *b*_*t−*1_ +∊_*t*_, where *∊*_*t*_ are independent with each other with mean 0 and variance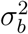. Here, in order to consider the trend of *b*_*t*_, we introduce the parameter *β*_*t*_ to be the trend slope of *b* from *b*_*t*_ to *b*_*t*+1_. The reason of doing this is that the model will contain a term *β*_*t*_ that explicitly represents the rate of change for a species at any given time. Therefore, this allows us to assess whether a species is “increasing”, “decreasing” or “stable” directly from the model output without any need for further processing. Then, the year effect is modelled by a pair of equations:

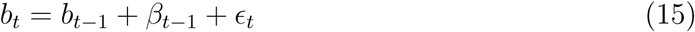

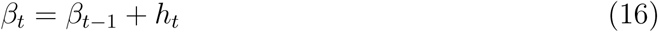

Here *∊*_*t*_ and *h*_*t*_ are all independent of each other with mean 0 and variance 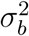 and 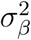 respectively. *β*_*t*_ is modelled by the random walk process. So equation 16 allows the slope *β*_*t*_ to change over time. Hence, *b*_*t*_ depends both on the previous one and the slope between them. This model is called local linear trend model in Chandler and Scott (2011). Then, according to this model, we update priors of the year effect *b*_*t*_ in equations 13 and 14 in the random walk model as follows:

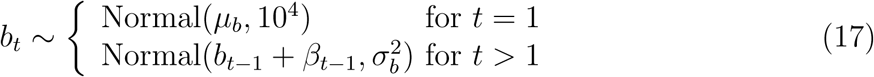

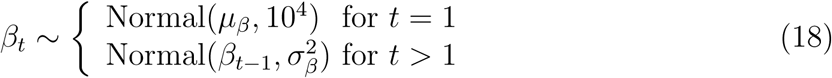

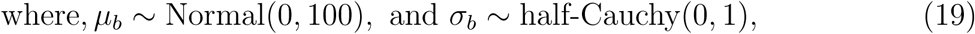

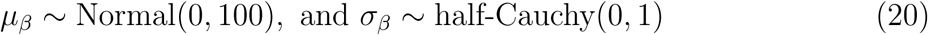

All other priors remain unchanged. Since we applied the local linear trend model to the prior, this occupancy model is referred to as the local linear trend model. Here the variance 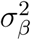 controls the variation of slope each year and both of the variance 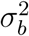 and the slope *β*_*t*_ allow the year effect *b*_*t*_ to vary over time. Therefore, the year effect *b*_*t*_ can have greater variation than that in the random walk model. Also, the estimation of slope will indicate the changes in year effect. For example, if the slope *β*_*t*_ is a positive number, then it means the proportion of occupied sites is expected to increase from year *t* to *t* + 1 and vice versa.

However, there is a potential problem in the local linear trend model. Although the introduction of the slope *β*_*t*_ allows greater changes in year effect *b*_*t*_, it may also causes violent fluctuations in the trend. Because the random walk process, just as its name, allows the value of *β*_*t*_ to increase or decrease randomly. Therefore, the slope can become infinite which is unrealistic in the natural world.

#### 3.1.2 Autoregressive local linear trend model

Because of the potential great fluctuations in the local linear trend model, we introduced a parameter *ϕ* whose absolute value is smaller than 1. This parameter may keep the variation of *b*_*t*_ within reasonable bounds. So the slope *β* is modelled by: *β*_*t*_ = *ϕβ*_*t−*1_ + *h*_*t*_ where *h*_*t*_ is still independent of each other with zero mean and constant variance. This is called first order autoregressive (AR(1)) model (Chandler and Scott, 2011). This model is stationary if and only if the absolute value of the parameter *ϕ* is strictly smaller than 1. Stationary means that the expectation and variance of *β*_*t*_ are equal for different *t* and the correlations between each pair of *β* at time points with same interval are equal, i.e. *corr*(*β*_*t*_, *β*_*t*+*k*_) = *corr*(*β*_*s*_, *β*_*s*+*k*_) (Chandler and Scott, 2011). The AR(1) process is stationary when the parameter is smaller than 1, while the local linear trend process is not stationary (with parameter equal to 1). Hence, the variation of *β*_*t*_ is smaller in the AR(1) process.

Next, we apply this process to the occupancy model. So equations 18 and 20 in the local linear trend model are replaced by the following:

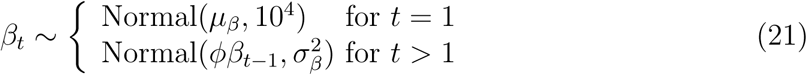

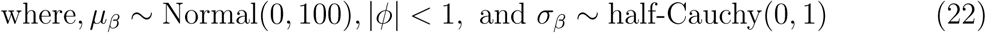

All other priors remain the same. Here we applied the stationary AR(1) model to the slope *β*_*t*_ and this model is based on the local linear trend model, so this occupancy model is referred to as autoregressive local linear trend model.

#### 3.1.3 Stationary autoregressive model

Since we applied AR(1) process to model the slope *β*_*t*_ in the last section, we try to apply this process to model the year effect *b*_*t*_ in random walk model. Hence, we replaced the equation 13 in the random walk model by the following:

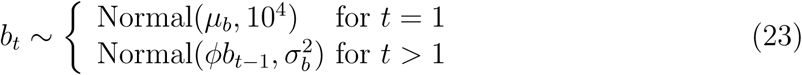

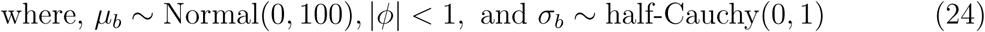

All other priors in the random walk model do not change. Here the AR(1) process is stationary with parameter *|ϕ| <* 1, so model is referred to as stationary autoregressive model.

### 3.2 Double random walk model

In Chapter 3.1, we discussed possible improvements of year effect *b*_*t*_. However, the prior of year effect *a*_*t*_ in equation 8 still has constant mean and variance. Hence, we applied the random walk process to model the year effect *a*_*t*_. We consider this separately from the models mentioned in section 3.1 because we are still not sure whether these models will be better than the random walk model or not. So the update is based on the random walk model by changing the equation 8 and 9 to the follows:

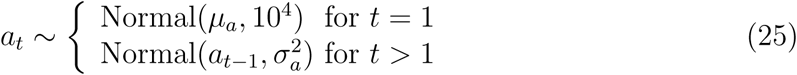

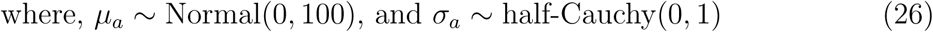

All other priors in the random walk model remain unchanged. Here, the mean of *a*_*t*_ depends on the previous one and *a*_*t*_ has constant variance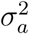. If the variance is large, the trend of *a*_*t*_ will vary greatly. Otherwise, if the variance is very small, the trend of *a*_*t*_ will be similar. This occupancy model is referred to as double random walk model because both of the year effects *b*_*t*_ and *a*_*t*_ are modelled by random walk process.

### 3.3 Spatial random field model

We have considered potential improvements in the year effects in Sections 3.1 and 3.2. Next, we will consider the prior of site effect *u*_*i*_ in equation 6 and 7 based on the random walk model.

The site effect *u*_*i*_ of site *i* may depend on the neighbourhood of this site. This means if the species is present in one site, it may also appear in the neighbour of this site. Hence, we use the Markov random field (MRF) model tested by Best et al. (2005) to model the distribution of site effect depending on other sites. The prior of MRF model specifies the distribution of *u*_*i*_ conditional on all of the other sites, as following:

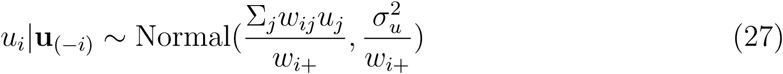

Here, **u**_(*−i*)_ denotes a vector of the site effects *u* in all sites except site *i*. *w*_*ij*_ is the *ij* th element of the matrix **W** with all diagonal entries *w*_*ii*_ = 0. The element *w*_*ij*_ = 1 if site *j* is the neighbour of site *i* and *w*_*ij*_ = 0 otherwise. Then the term *w*_*i*+_ is the total number of neighbours of site *i*. Since the locations in the dataset are recorded on a 1 *km*^2^ grid, it is common to define the neighbours of any grid cell to be the other cells that are horizontally or vertically adjacent.

Here, we need at least 90,000 sites to fit model using the hoverfly dataset. However, the software we used to fit this model cannot work with so many sites. Hence, we combine every 100 sites to form a big square site. Therefore, the area of each site changes from 1*×*1 *km*^2^ to 10*×*10 *km*^2^. Besides, the software is prone to errors if the distribution of standard deviation *σ* is half-Cauchy(0,1). As a result, in this model, we change the distribution of all standard deviations *σ* from half-Cauchy back to uniform distribution. Hence we replace all priors in random walk model with the follows:

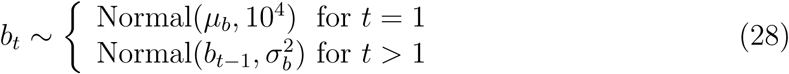

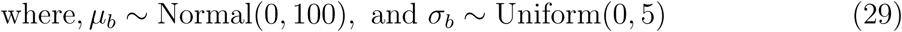

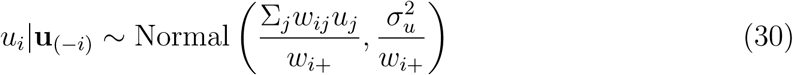

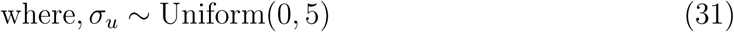

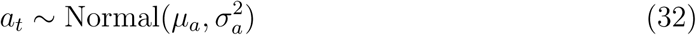

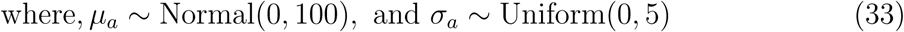

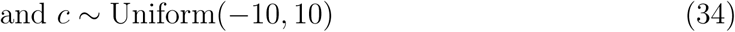

This model aims to find a better model in the prior of site effect *u*_*i*_ using Markov random field model, so this occupancy model is referred to as spatial random field model.

### 3.4 Categorical list length model

The final part we consider to improve in this research is the parameter *c* which represents the relationship between sampling effort and the detection probability. In the base model, the logit of the detection probability has a logarithmic relationship with the list length. However, this has not been verified. Hence, in this section, we will shift the parameter *c* to a categorical variable which has different values with different list lengths. Here, we replaced *c log*(*L*_*itv*_) in equation 4 with 21 categories as follows:

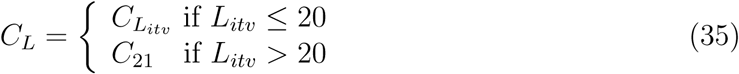

Here, we choose 20 as a boundary of the list length in the variable *C*_*L*_ since over 98% of the list lengths are not longer than 20 in the dataset. Next, we applied this variable into the detection submodel in equation 4 as following:

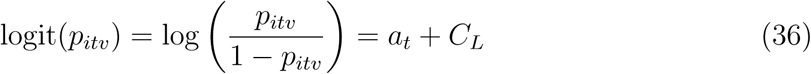

As a result, the prior of *c* in equation 10 is replaced by the prior of variable *C* as following:

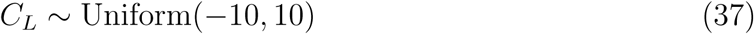

This model is referred to as the categorical list length model. After fitting the model, we can plot the relationship between variable *C*_*L*_ and list length *L* to see whether the detection probability has a logarithm increasing when the sampling effort increases. Further details about the result of this model will be shown in Chapter 5.

### 3.5 Summary

This chapter aims to suggest some improvements to the random walk model mentioned in Chapter 2 built by Outhwaite et al. (2018). Based on the structure of occupancy-detection model in equation 3 and 4 in Chapter 2.1, the potential improvements contain four parts, which are priors of two year effects and the site effect, and the relationship between list length and detection probability. The code of the model definition of each model is provided in the appendix. Further information about fitting these models are explained in Chapter 4 and the results of these models are shown in Chapter 5.

## 4 Model fitting

The previous two chapters reviewed some occupancy-detection models that have been discussed in the literature, and suggested some potential improvements to these models. This chapter discusses the methods that are used to fit these models in practice. As discussed in Chapter 2, the models have complex structures and many unknown parameters, and Bayesian computational techniques provide a convenient way to fit such models. We start by introducing these computational techniques. Then we show the software and packages used to fit each model and tools of comparing the models. Finally, the process of choosing initial values for each variables in each model will be provided.

### 4.1 Markov chain Monte Carlo (MCMC) method

As mentioned in Section 2.1, we use Bayesian methods to estimate parameters. To find the posterior distribution of the parameter, we use the iterative methods known as Markov chain Monte Carlo (MCMC) (Lunn et al., 2013). The iterative sampling scheme is as follows: (i) choosing the initial values of the parameter vector ***θ***^(0)^, (ii) sampling the new values of ***θ***^(*k*)^ from a suitably-chosen distribution conditional on the previous values ***θ***^(*k−*1)^, (iii) repeating the step 2 many times. Finally the parameter will sample from the posterior distribution. The whole process is treated as one chain and the number of repetitions of step 2 is known as the number of iterations.

Since MCMC samples from posterior distribution eventually, we need to discard the initial iterations which are not sampling from the posterior distribution. The discarded period is called ‘burn-in’ period. To identify this burn-in period, we can run multiple chains with different initial values. In this research, we will run 3 chains for each model. Then the burn-in period will be the period before all these chains converge to the same region. One of the formal way to detect the convergence is to observe the value of Gelman-Rubin statistic (often known as 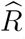,) of each variable. The ratio 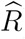, is calculated using the formula: 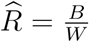, where *B* is the width of the 100(1-*α*)% credible interval taken from all saved iterations and *W* is the width of the 100(1-*α*)% credible interval calculated from the posterior distribution with parameter from the final iteration (Lunn et al., 2013). This value will always be greater than or equal to 1 and should theoretically be equal to 1 if all of the chains are sampling the same distribution. In this research, we assume convergence if 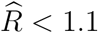 for all variables, which is a standard choice in many literatures(Melinscaka and Montesanoa, 2016; Farquharson et al., 2018).

However, there is a potential difficulty of the MCMC method that the consecutive samples may be correlated. This situation is called autocorrelation. To reduce the autocorrelation, we keep only every 10th value in this research to make these samples approximately independent. This process is called ‘thinning’.

### 4.2 Tools for fitting the model

The software we use in this research to fit the models are JAGS (Plummer, 2003) and OpenBUGS (Spiegelhalter et al., 2014). Both of these two software are designed for Bayesian analysis using MCMC techniques. In most situations, we will use JAGS to fit the model since JAGS has a shorter running time than OpenBUGS. Also, JAGS allows the distribution of standard deviation *σ* to be half-Cauchy(0, 1) as shown in equations 7 and 9 while OpenBUGS keeps getting errors in this situation. This is because JAGS and OpenBUGS have different updating strategies, and the updating strategy in OpenBUGS is prone to errors with the half-Cauchy distributions. However, JAGS cannot fit the spatial random field model in Section 3.3. Hence, we will use OpenBUGS to fit this model. That is the reason why we change the distribution of standard deviation *σ* from half-Cauchy to Uniform distribution in Section 3.3.

Since we will also use R (R Core Team, 2015) to handle data and make plots, we use the package R2JAGS (Su and Yajima, 2015) and R2OpenBUGS (Sturtz et al., 2005) to run JAGS and OpenBUGS through R. The R codes of the model definitions are provided in the appendix.

### 4.3 Model comparison

After fitting the model, some tools are needed to compare the performance of each model. Therefore, in this section, we introduce the tool will be used to compare the models in Chapter 5, which is called Deviance Information Criterion (DIC). DIC is a tool to compare the complexity and fit of models (Spiegelhalter et al., 2002). The formula of DIC of the parameter is: DIC = 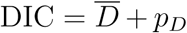 + *p*_*D*_, Where, 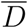 is the posterior mean deviance which is minus twice the log-likelihood of the posterior distribution. This part reflects the model fit. *p*_*D*_ can be treated as a penalty term of the model complexity (Best et al., 2005). If the model is complex, the value of *p*_*D*_ will be high. Therefore, the model will be better if the value of DIC is lower.

### 4.4 Initial values

As stated in Section 4.1, the initial values *θ*^(0)^ should be set in the first step to start the MCMC iterations. Although the software can generate the initial values automatically, we will choose them manually. Because the initial values of variables should be set carefully so that the model will converge quickly with small number of iterations. The initial values need to be sufficiently widely dispersed that the use of multiple chains provides some reassurance about convergence, but they should not be in areas with extremely low posterior probability because this can lead to very long convergence times and also to numerical problems associated with the calculation of ratios of very small quantities. Hence, in this section, we introduce how to choose the initial values of each parameter in different models. Each subsection below will introduce the way of choosing the initial values of all parameters related to one type of unknown quantities.

#### 4.4.1 The year effects

Firstly, we can set the initial values of year effects *a*_*t*_ and *b*_*t*_ based on the proportion of detection for each species which is the ratio of the number of detections and the number of visits. We calculate the overall detection proportion and also the proportions for each year. Then the logit of these proportions are calculated using the formula: logit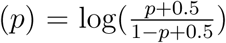, where *p* is the proportion of detection. Here we add 0.5 on both numerator and denominator to make sure the value inside the logarithm will not be so close to 0 or 1 that the logarithm will tends to minus or plus infinity. This transformation is known as empirical logit transform (Agresti, 1990).

Next, we can calculate the standard deviation, *σ*, of the logit of a probability *p* estimated from *n* independent binary variables based on the propagation of uncertainty formula. Then the initial values of *a*_*t*_ are set as the sum of the logit detection proportion each year and the random values from normal distribution with mean 0 and standard deviation equal to 4*σ*. Similarly, the initial values of *b*_*t*_ are set to be negatively correlated to those of *a*_*t*_. So the initial values of *b*_*t*_ are logit detection proportion each year minus the random values from normal distribution with mean 0 and standard deviation 4*σ*.

Then, the initial values of the standard deviations of *a*_*t*_ and *b*_*t*_ can be considered as the standard deviations of the initial values of the year effects *a*_*t*_ and *b*_*t*_. The reason why these choices might be appropriate is that the initial values of year effects combine the simulations from normal distribution and the empirical values from dataset. So the initial values of the standard deviations are based on the dataset, but will not be influenced much by the special cases in the dataset. All these initial values are applied to all of the models except the local linear trend model and autoregressive local linear trend model as shown in Sections 3.1.1 and 3.1.2, because the year effect *b*_*t*_ of these two models are based on the slope β_*t*_.

Therefore, we need to set the initial values of *a*_*t*_, *b*_*t*_ and β_*t*_ separately for these two models. The initial values of β_*t*_ are considered first. The initial values of β_*t*_ are random from the normal distribution as before. Then the initial values of *b*_*t*_ are set to be the sum of the initial values of β_*t*_ + *b*_*t−*1_ and the random values from the normal distribution with mean 0 and standard deviation 2*σ* if t*≥*2. The reason of setting a smaller standard deviation here is that the initial values of β_*t*_ have already got the random part with a standard deviation as usual, so the initial values of *b*_*t*_, which are the summation of *β*_*t*_ and *b*_*t−*1_, already have the random part from the initial values of *β*_*t*_. So intuitively, the initial values of *b*_*t*_ only need the random part with a smaller standard deviation. Then the initial value of *b*_1_ is chosen to be the sum of logit detection proportion for the first year and the random value from normal distribution with mean 0 and standard deviation 4*σ* as before. The reason of setting the initial values of *b*_*t*_ and *β*_*t*_ like these is that *β*_*t*_ is the difference between two consecutive values of *b*_*t*_, if we choose the initial values of *b*_*t*_ and *β*_*t*_ randomly, it will not match the relationship between these two quantities. Then, the initial values of *a*_*t*_ are chosen as the negative initial values of *b*_*t*_. In addition, we set the initial values of the standard deviations of *a*_*t*_, *b*_*t*_ and *β*_*t*_ are the standard deviations of their initial values.

#### 4.4.2 The site effect

We use a similar approach to choose the initial values of the site effect *u*_*i*_. Firstly, the logit proportion of detection for each site is calculated based on the dataset for each species. Then the initial values of *u*_*i*_ are the sum of the detection proportion and the random values from the normal distribution with mean 0 and standard deviation 4*σ*, where *σ* is equal to the standard deviation of the logit one as shown in Section 4.4.1. Hence, the initial values of the standard deviation of *u*_*i*_ is set to be the standard deviation of its initial value.

However, the initial values of the site effect *u*_*i*_ for spatial random field model should be considered individually. OpenBUGS will be used to fit this model as explained in Section 4.2. But OpenBUGS will work for this model only if the mean of initial values of *u*_*i*_ is exactly 0. Therefore, we set all initial values of *u*_*i*_ to be 0. But we cannot set the initial values of the standard deviation of *u*_*i*_ to be zero and the spatial model is prone to convergence problems unless the initial values and priors for standard deviations are chosen very carefully. So the results from other models are used to inform the settings here. Therefore, we set the initial value of standard deviation of *u*_*i*_ randomly from Uniform(0.7, 1.1) because usually the initial values of the standard deviation of *u*_*i*_ in other models are distributed from 0.7 to 1.1.

#### 4.4.3 The parameter of list length

Next, we choose the initial values of the parameter of the list length *c* for all models excluding the categorical list length model. Notice that from equation 4, the logit of the probability of detection has a linear relationship with the logarithm of the list length. To determine some plausible starting values therefore, we carry out a logistic regression of the observed detections on the logarithm of the list length *L*_*itv*_, and randomly sample the initial value of *c* from a normal distribution centred on the coefficient of log *L*_*itv*_ in this fitted model, and with variance equal to twice the reported variance of the coefficient estimate.

However, we need to consider the categorical list length model separately. We use JAGS to fit this model as mentioned in Section 4.2. JAGS will work only if there is no self-chosen initial values for the categorical vector *C*_*L*_, and then JAGS will generate initial values for *C*_*L*_ itself. Therefore, we do not set initial values of *C*_*L*_ in this model.

#### 4.4.4 True occupancy status

In addition, we will set the initial values of the true occupancy status *z*_*it*_. For cases where the focal species was observed, clearly *z*_*it*_ must be 1. For other cases, initial values of the true occupancy status *z*_*it*_ were sampled from a Bernoulli distribution with parameter 0.5.

### 4.5 Summary

This chapter first introduces the MCMC method to be used in Bayesian analysis to fit the models, then explains the two software of fitting the model using MCMC method, which are JAGS and OpenBUGS. Also, the packages are introduced to run JAGS and OpenBUGS through R. Then, this chapter provides a tool to compare the models which is called DIC. Finally, this chapter describes the procedure of choosing initial values for each variable in each model.

## 5 Comparison of model performance

In previous chapters, we reviewed some occupancy-detection models, suggested some potential improvements to the models and explained the methods used to fit these models in practice. In this chapter, we use the dataset of anasimyia contracta and anasimyia interpuncta introduced in Section 1.2 to fit all occupancy-detection models in Chapters 2 and 3. For each species, the analyses in this chapter are confined to the subset of sites where this species is neither always observed nor never observed: we refer to these as ‘informative sites’. The reason of doing this is to speed up the model fitting process by not wasting time over the majority of sites where the species is never observed in the past. In this chapter, we will first compare the MCMC convergence of each model based on their convergence situations, the number of iterations and the time used for a model to converge. Then we will show the plots of the trends of the species anasimyia contracta and anasimyia interpuncta using these occupancy models. Finally, there will be a comparison of the complexity and fit of each model using the values of DIC as explained in Section 4.3.

### 5.1 Convergence

We use the software introduced in Section 4.2 to fit models with initial values set in Section 4.4. The numbers of iterations of each model and the time of running these models for the species anasimyia contracta and anasimyia interpuncta are shown in Table 1 and 2 respectively in order to compare the speed of convergence of the models. All running times are based on the laptop MacBook Air with Windows operation system and the processor speed is around 800 MHZ. Also, the values of 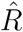are examined to determine whether the model is convergent or not.

**Table 1:** The tables shows the numbers of MCMC iterations and burn-in, the running time and the software used to fit all models with anasimyia contracta dataset.

| Models | Iterations | Burn-in | Running time | Software |
| --- | --- | --- | --- | --- |
| The base model | 10,000 | 5,000 | 45min | JAGS |
| Adaptive stationary model | 10,000 | 5,000 | 45min | JAGS |
| Random walk model | 20,000 | 10,000 | 1.5h | JAGS |
| Local linear trend model | 25,000 | 15,000 | 2h | JAGS |
| Autoregressive local linear trend model | 25,000 | 15,000 | 2h | JAGS |
| Stationary autoregressive model | 20,000 | 10,000 | 1.5h | JAGS |
| Double random walk model | 25,000 | 15,000 | 2h | JAGS |
| Spatial random field model | 5,000 | 3,000 | 34h | OpenBUGS |
| Categorical list length model | 20,000 | 10,000 | 1.5h | JAGS |

**Table 2:** The table shows the numbers of MCMC iterations and burn-in, the running time and the software used to fit all models with anasimyia interpuncta dataset.

| Models | Iterations | Burn-in | Running time | Software |
| --- | --- | --- | --- | --- |
| The base model | 20,000 | 10,000 | 5min | JAGS |
| Adaptive stationary model | 70,000 | 50,000 | 14min | JAGS |
| Random walk model | 150,000 | 130,000 | 40min | JAGS |
| Local linear trend model | 250,000 | 230,000 | 1h | JAGS |
| Autoregressive local linear trend model | 250,000 | 230,000 | 1h | JAGS |
| Stationary autoregressive model | 200,000 | 180,000 | 47min | JAGS |
| Double random walk model | 250,000 | 230,000 | 1h | JAGS |
| Spatial random field model | 20,000 | 10,000 | 7.5h | OpenBUGS |
| Categorical list length model | 20,000 | 10,000 | 1.5h | JAGS |

All models in Table 1 and 2 are run for three Markov chains with a thinning rate of ten. From the number of iterations in the table we can see that the MCMC of the base model uses the least time to converge with the smallest number of iterations for both species. Then follows the adaptive stationary model and the random walk model. The categorical list length model uses more time to be fitted than the random walk model. However, the local linear trend model, the MCMC of the autoregressive local linear trend model and the double random walk model are the most difficult to converge. Although they are run for 250,000 iterations for anasimyia interpuncta, they still do not converge with the values of 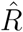for many parameters are much greater than 1.1. The values of 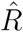are even greater than 3 for some parameters. Therefore, these three models are not suitable for low recording intensity species. Moreover, the MCMC of the autoregressive local linear trend model is the most difficult model to converge: for this model, some parameters still do not converge for anasimyia contracta while other models for this species all converge with the same number of iterations. Finally, the spatial random field model is fitted using OpenBUGS which has different updating system to JAGS. Hence, the spatial random field model converges with small number of iterations in a longer time. Each iteration of the MCMC for this model is very slow for anasimyia contracta due to the huge dataset, therefore it will need several days to run more iterations. Although convergence was achieved for most parameters, it it not possible to converge for all of the parameters with 5,000 iterations. We can also conclude that the spatial random field model is really time-consuming for large dataset with many records.

### 5.2 Performance of each model

The results of each model fitting the anasimyia contracta and anasimyia interpuncta data will be shown in this section. Each plot shows the proportion of occupied sites with 95% credible intervals. The solid lines in the plots show the posterior means for the proportions, whereas the shaded parts show the 95% credible intervals. The proportion of occupied sites is calculated using the number of sites the species occupied divided by the number of informative sites. This is why the proportions in the plots are very high although these two species are relatively scarce.

#### The base model

The results of the base model with dataset anasimyia contracta and anasimyia interpuncta are shown in Figure 3. The trend of the proportion fluctuates violently from year to year which is not realistic in the nature. Also, the credible interval is so wide that the values of credible intervals are from 0 to 1 in some years which means the base model is not very precise. In addition, the fluctuation is greater and the credible interval is wider in the plot of the anasimyia interpuncta. This is because this species has fewer records, with only 79 records in the past 45 years as shown in Figure 1.

**Figure 3:**
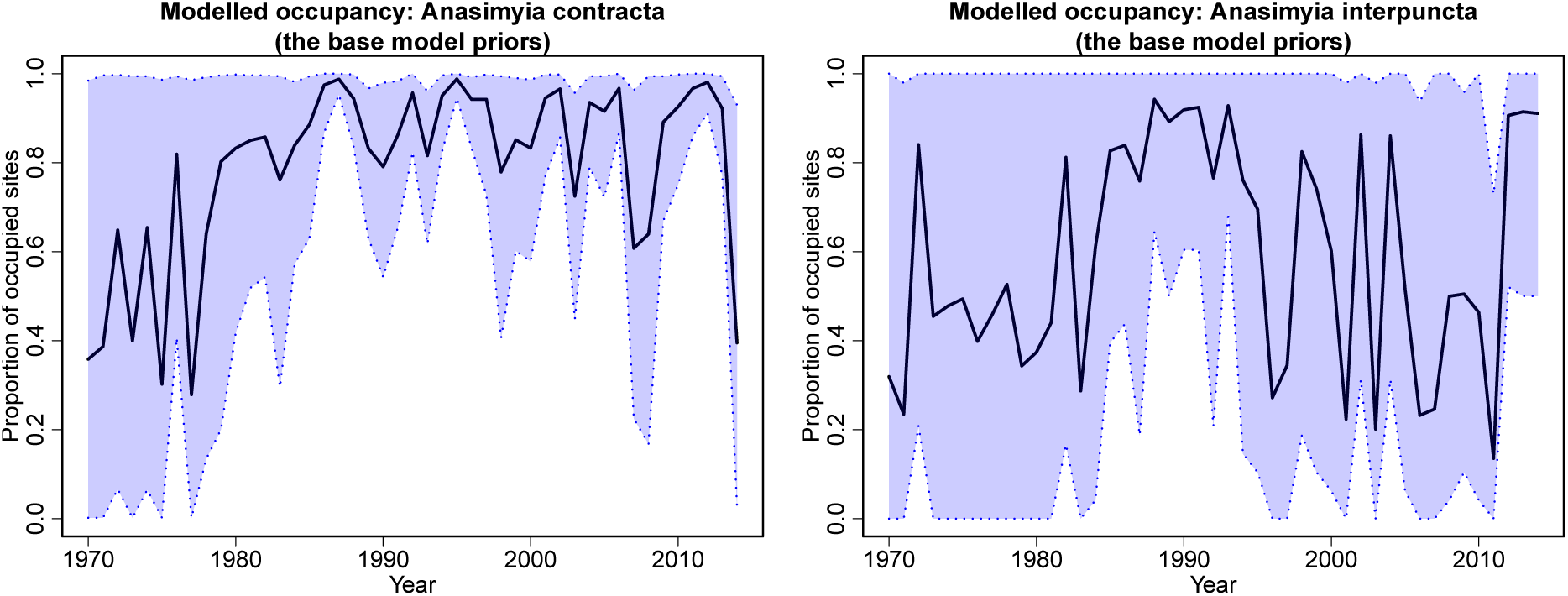
The solid black lines show the posterior means of the proportions of occupied sites using the base model with only informative sites from 1970 to 2014. The blue shaded areas show the 95% credible interval of the proportion. The left hand side plot uses the dataset of anasimyia contracta and the right hand site plot uses the dataset of anasimyia interpuncta.

#### Adaptive stationary model

The proportions of occupied sites for anasimyia contracta and interpuncta with informative sites are shown in Figure 4. The trends are smoother and the credible intervals are narrower than those of the base model. Moreover, the credible intervals are narrow at the start and end of the time period when there are fewer records. However, the result of the simulations experiment by Outhwaite et al. (2018) shows that the adaptive stationary model is the most biased model among three models. Therefore, having narrow credible interval for this unjustified model is dangerous since the real proportion of occupied sites may not be in the range of the credible interval. Actually, the simulations experiment also shows that the adaptive stationary model has the lowest coverage rate among all three models presented in Outhwaite et al. (2018). Hence, the credible intervals of the adaptive stationary model are too narrow to cover the 95% of the actual occupancy proportions.

**Figure 4:**
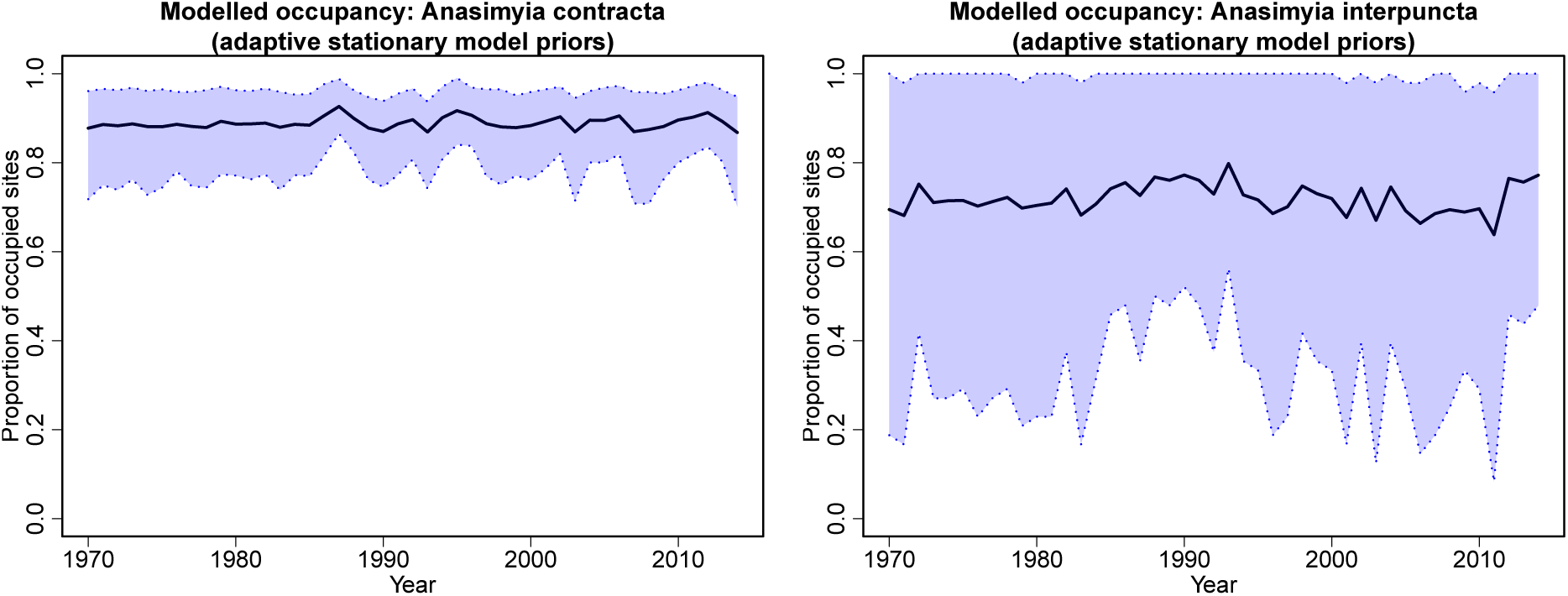
The solid black lines show the posterior means of the proportions of occupied sites using the adaptive stationary model with only informative sites from 1970 to 2014. The blue shaded areas show the 95% credible interval of the proportion. The left hand side plot uses the dataset of anasimyia contracta and the right hand site plot uses the dataset of anasimyia interpuncta.

Furthermore, the trend of the species still goes up and down frequently from year to year, although the magnitude is smaller than the base model. Also, the year effect *b*_*t*_ has a constant mean as shown in Section 2.2 which restricts the greater changes of theproportion of occupied sites. That is why the overall trend of each species is flat.

#### Random walk model

The trends of the proportions of occupied sites for anasimyia contracta and anasimyia interpuncta with 95% credible interval by the random walk model is shown in Figure 5. The trend fluctuates little over time and the random walk model allows greater changes in the trend than the adaptive stationary model. The credible intervals of the random walk model with more records are narrower which means the random walk model is more precise than the previous two models if the number of records is large. However, if the records are fewer, the credible intervals of the random walk model is wide because the uncertainty of the estimation is larger with smaller number of records. Although the credible interval of the random walk model is wider than that of the adaptive stationary model when there are fewer records, it is more realistic. As mentioned in the previous subsection, the coverage of the adaptive stationary model credible interval is much lower than 95% of the actual occupancy proportion. Therefore, the random walk model is moreprecise than the previous two models.

**Figure 5:**
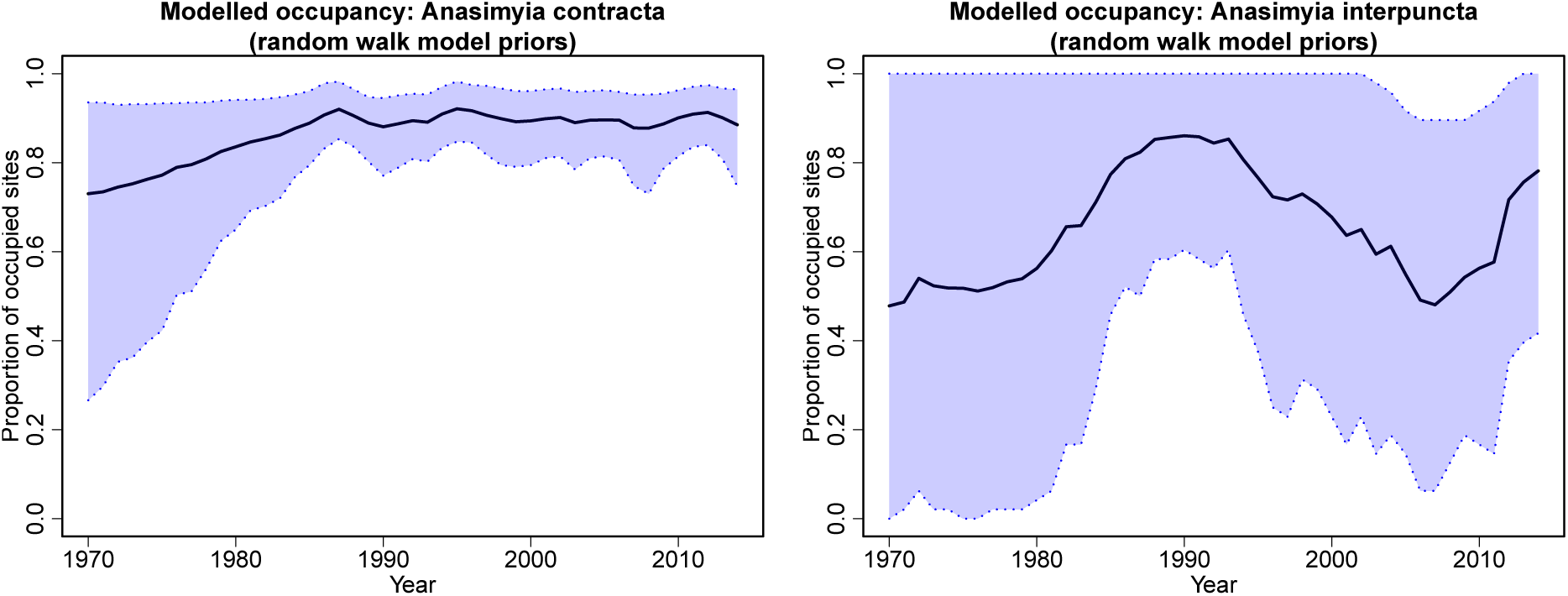
The solid black lines show the posterior means of the proportions of occupied sites using the random walk model with only informative sites from 1970 to 2014. The blue shaded areas show the 95% credible interval of the proportion. The left hand side plot uses the dataset of anasimyia contracta and the right hand site plot uses the dataset of anasimyia interpuncta.

#### Local linear trend model

The proportions of occupied sites for anasimyia contracta and anasimyia interpuncta with their 95% credible intervals using the local linear trend model are shown in Figure 6. The trend is smooth, however, the credible interval is wide when the records of the species are few.

**Figure 6:**
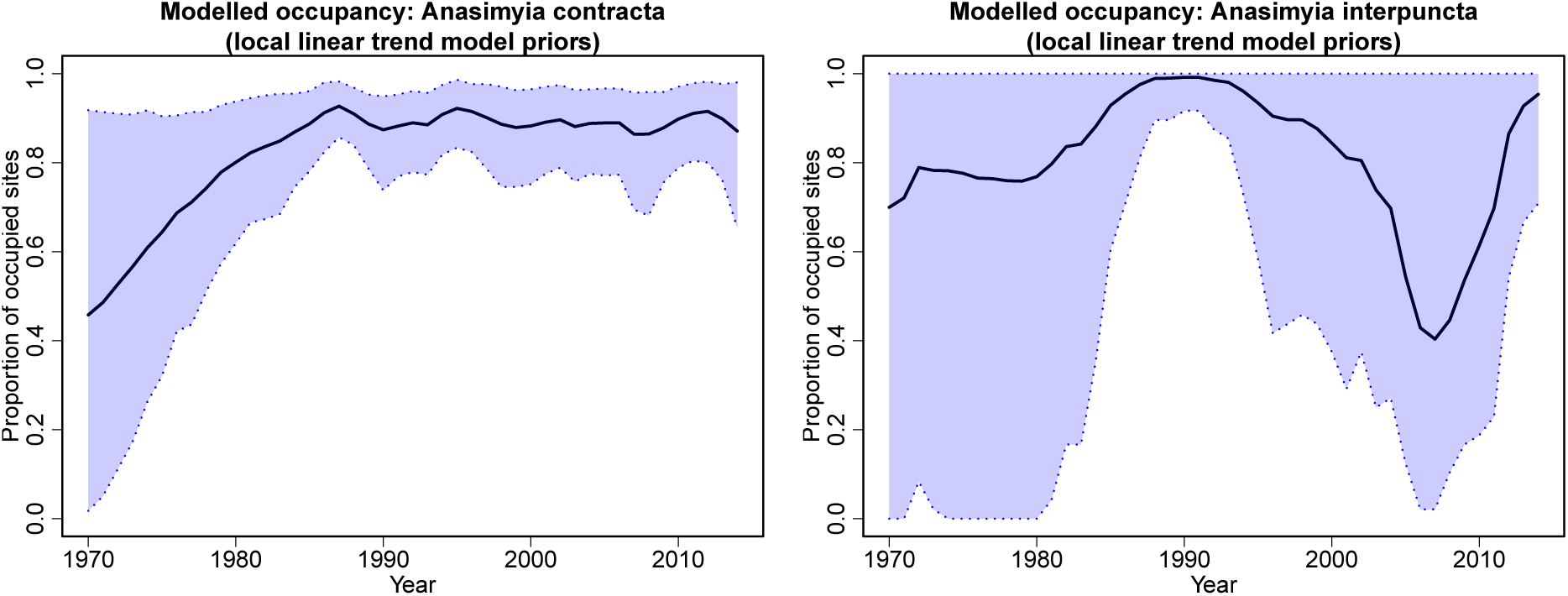
The solid black lines show the posterior means of the proportions of occupied sites using the local linear trend model with only informative sites from 1970 to 2014. The blue shaded areas show the 95% credible interval of the proportion. The left hand side plot uses the dataset of anasimyia contracta and the right hand site plot uses the dataset of anasimyia interpuncta.

As mentioned in Section 3.1.1, the reason of introducing the slope is that the parameter *β*_*t*_ can show the trend of occupied sites directly. For anasimyia contracta, most of the trends in the plot match the value of the *β*_*t*_. However, there is a decrease from 1987 to 1990 of the solid line in the plot of anasimyia contracta, while the values of *β*_*t*_ are 0.071, 0.033 and 0.015 respectively. Here the value of *β*_*t*_ is the mean of the posterior distribution of *β*_*t*_ and the solid line shows the posterior mean of the proportion of occupied sites. But the actual values of *β*_*t*_ and the proportion of occupied sites may not be the mean of their posterior distributions. So there may be some uncertainties of these posterior means.

Since the posterior means of *β*_*t*_ are very close to zero, although the trend of the species is downward, the posterior mean of *β*_*t*_ can still be positive in the same period.

However, things are different for anasimyia interpuncta. There is a significant increase from 1980 to 1987 in the plot of anasimyia interpuncta. However, the mean values of *β*_*t*_ are all negative which do not show any signal of increasing. Actually, the local linear trend model for anasimyia interpuncta is really difficult to converge which was shown in Section 5.1 and the MCMC for this model still has not converged although the initial values are set really carefully as explained in Section 4.4. This lack of convergence seems to be the most likely explanation for the apparent incompatibility between the underlying trend and the estimates of *β*_*t*_ for this species.

#### Autoregressive local linear trend model

The proportions of occupied sites for anasimyia contracta and anasimyia interpuncta with their 95% credible intervals using the autoregressive local linear trend model are shown in Figure 7. The trend is less smooth than that from the local linear trend model.Also, the credible interval is wide when the records of the species are few.

**Figure 7:**
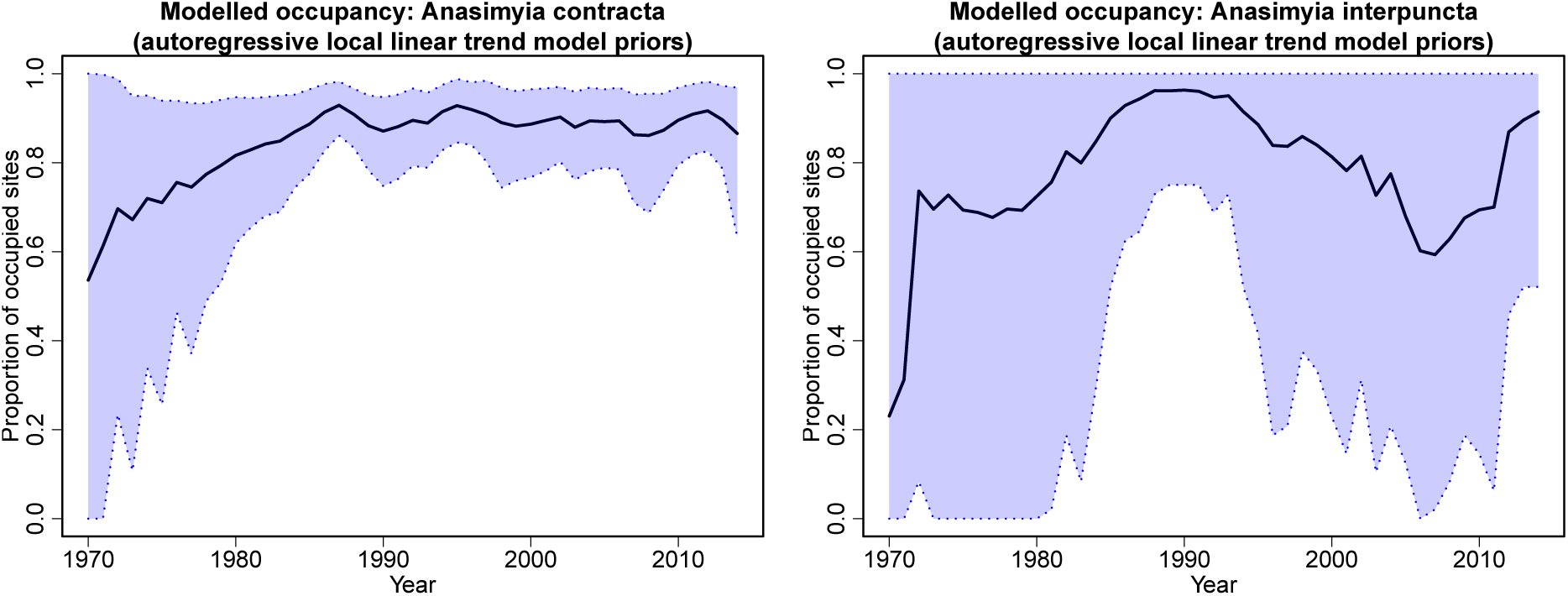
The solid black lines show the posterior means of the proportions of occupied sites using the autoregressive local linear trend model with only informative sites from 1970 to 2014. The blue shaded areas show the 95% credible interval of the proportion. The left hand side plot uses the dataset of anasimyia contracta and the right hand site plot uses the dataset of anasimyia interpuncta.

As mentioned in Section 5.1, the autoregressive local linear trend model is influenced a lot by the low recording intensity, so the MCMC for this model still has not converged for both species. Therefore, the posterior distribution of this model may not include the actual value of the parameter. Although the introducing of the slope *β*_*t*_ is to show the trend of the proportion of occupied sites directly, the lack of convergence may be the potential reason why the posterior means of *β*_*t*_ is not consistent with the trend of the proportion.

#### Stationary autoregressive model

The proportions of occupied sites for anasimyia contracta and anasimyia interpuncta with their 95% credible intervals using the stationary autoregressive model are shown in Figure 8. The trends fluctuate more violently than the random walk model and the adaptive stationary model and the credible interval is wide if the records are few. Besides, the overall trend is flatter than that of the random walk model since the stationaryautoregressive model does not allow the trend to increase or decrease greatly over time as the random walk model does.

**Figure 8:**
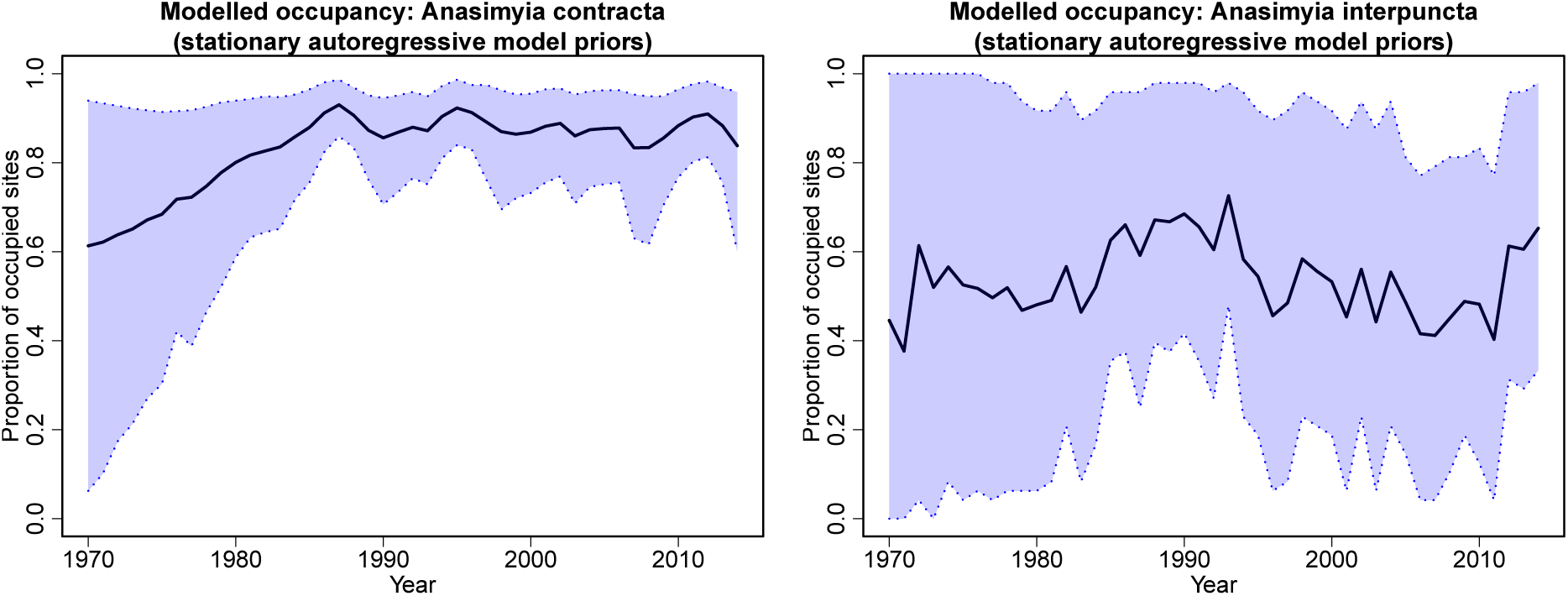
The solid black lines show the posterior means of the proportions of occupied sites using the stationary autoregressive model with only informative sites from 1970 to 2014. The blue shaded areas show the 95% credible interval of the proportion. The left hand side plot uses the dataset of anasimyia contracta and the right hand site plot uses the dataset of anasimyia interpuncta.

#### Double random walk model

The proportions of occupied sites for anasimyia contracta and anasimyia interpuncta with their 95% credible intervals using the double random walk model are shown in Figure The trend of anasimyia interpuncta is really flat while the trend of anaisimyia contracta does not show much difference with the previous models. Also, the credible intervals are wider than that of the random walk model.

#### Spatial random field model

The proportions of occupied sites for anasimyia contracta and anasimyia interpuncta with their 95% credible intervals using the spatial random field model are shown in Figure The trend of the spatial random field model is similar to that of the random walk model although the overall trend of the spatial random field model is flatter. In addition,the credible intervals are a little bit wider. Therefore, the addition of the dependence of the sites does not have great effect on the random walk model.

#### Categorical list length model

Finally, the proportions of occupied sites for anasimyia contracta and anasimyia interpuncta with their 95% credible intervals using the categorical list length model are shown in Figure 11. The trends for two species with this model are similar to those from the random walk model. However, there are greater fluctuations between some years while the overall proportions with the categorical list length model are lower than those with the random walk model. Besides, the credible intervals is narrower with the categorical list length model, but we cannot say that the precision of the categorical list length model is higher than that of the random walk model only from the credible intervals, since the we do not know the coverage of the credible interval of the categorical list length model.

**Figure 9:**
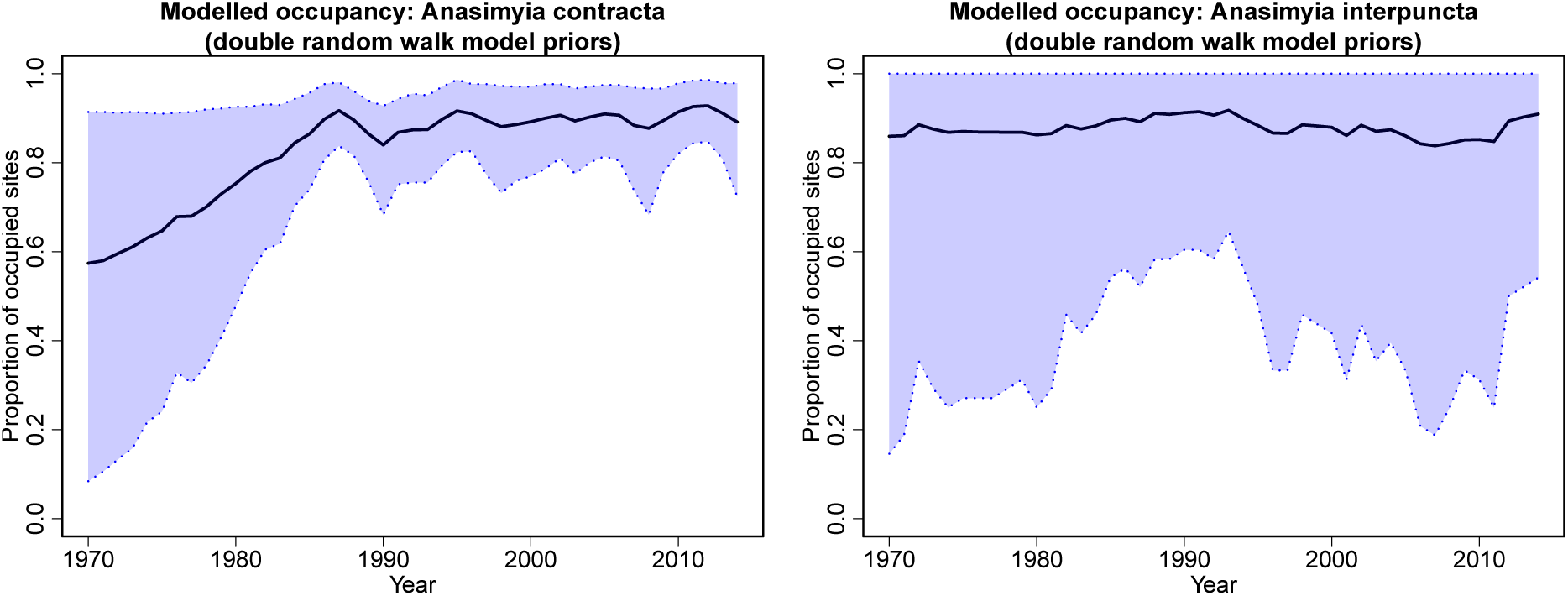
The solid black lines show the posterior means of the proportions of occupied sites using the double random walk model with only informative sites from 1970 to 2014. The blue shaded areas show the 95% credible interval of the proportion. The left hand side plot uses the dataset of anasimyia contracta and the right hand site plot uses the dataset of anasimyia interpuncta.

**Figure 10:**
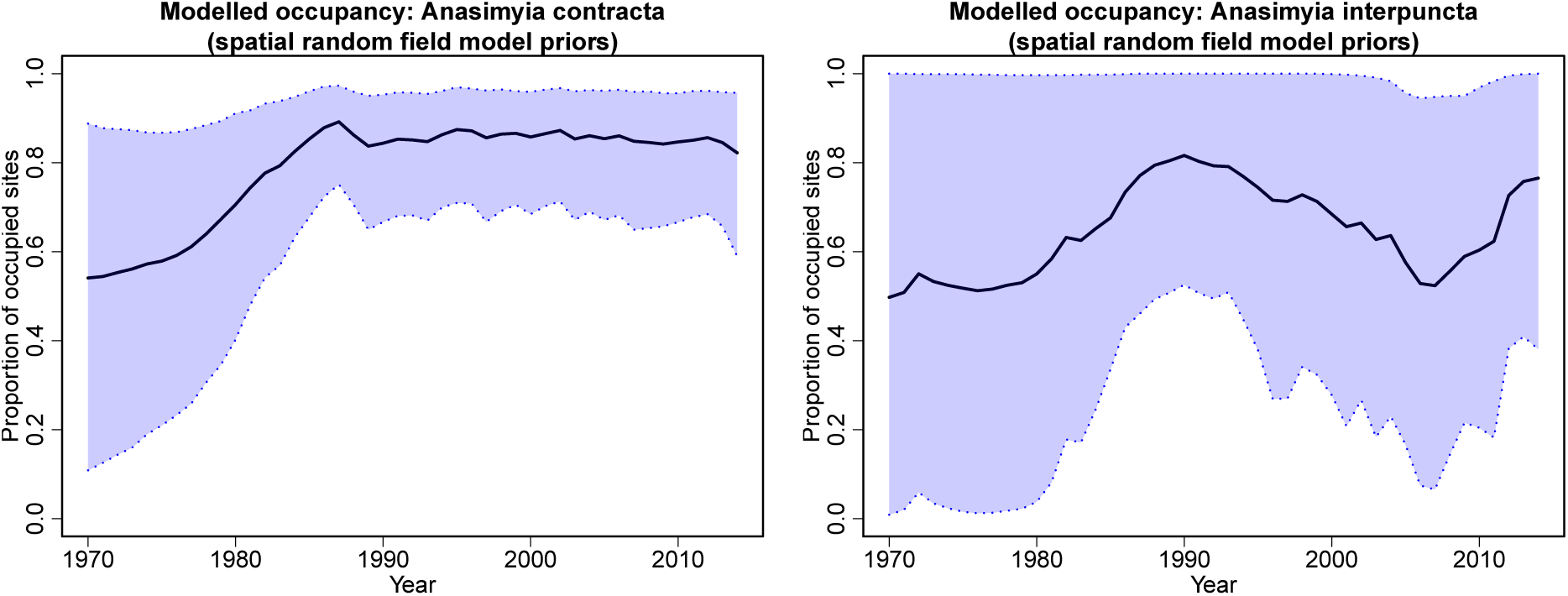
The solid black lines show the posterior means of the proportions of occupied sites using the spatial random field model with only informative sites from 1970 to 2014. The blue shaded areas show the 95% credible interval of the proportion. The left hand side plot uses the dataset of anasimyia contracta and the right hand site plot uses the dataset of anasimyia interpuncta.

**Figure 11:**
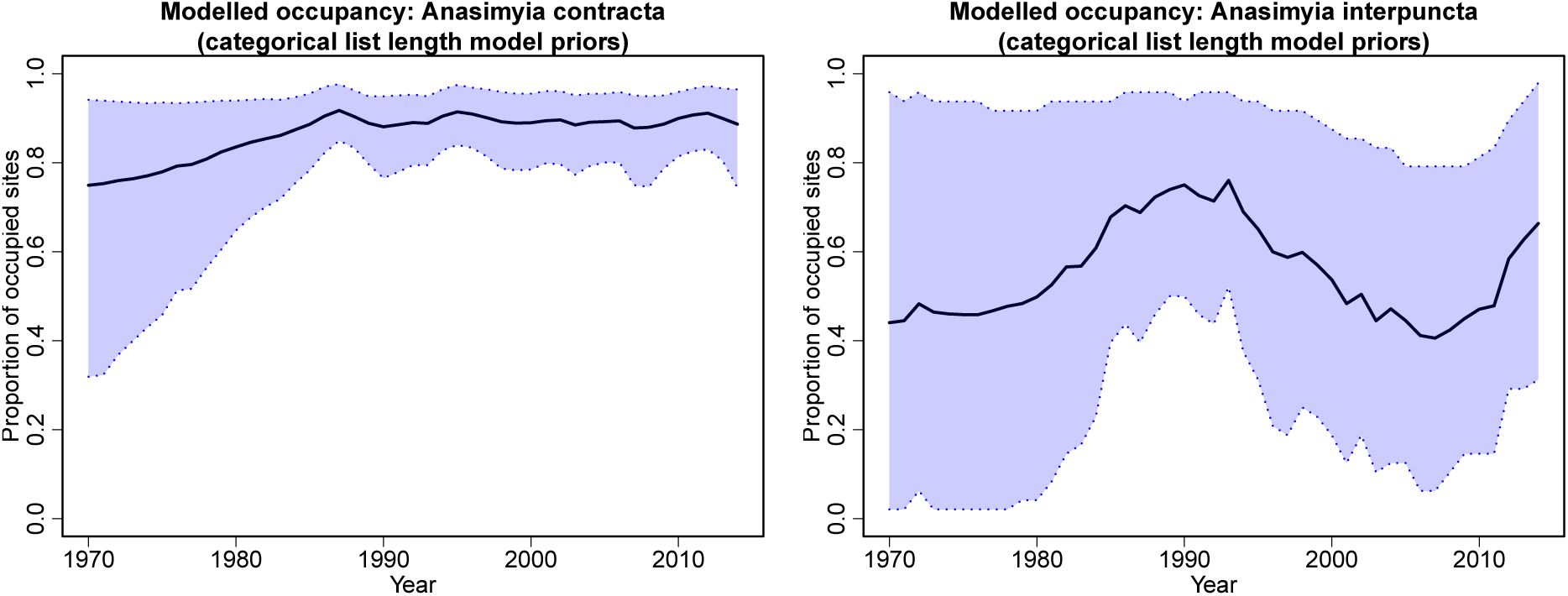
The solid black lines show the posterior means of the proportions of occupied sites using the categorical list length model with only informative sites from 1970 to 2014. The blue shaded areas show the 95% credible interval of the proportion. The left hand side plot uses the dataset of anasimyia contracta and the right hand site plot uses the dataset of anasimyia interpuncta.

The aim of introducing this model was to check the relationship between the list length and the probability of detection. There is a logarithmic relationship between thelist length and the probability of detection on the logit scale in the previous models as shown in equation 4. To verify this relationship, the plots between the list length and the categorical variable *C*_*L*_ are produced in Figure 12 together with the 95% credible intervals. This plot can show the relationship between list length and the probability of detection, as shown in equation 36.

**Figure 12:**
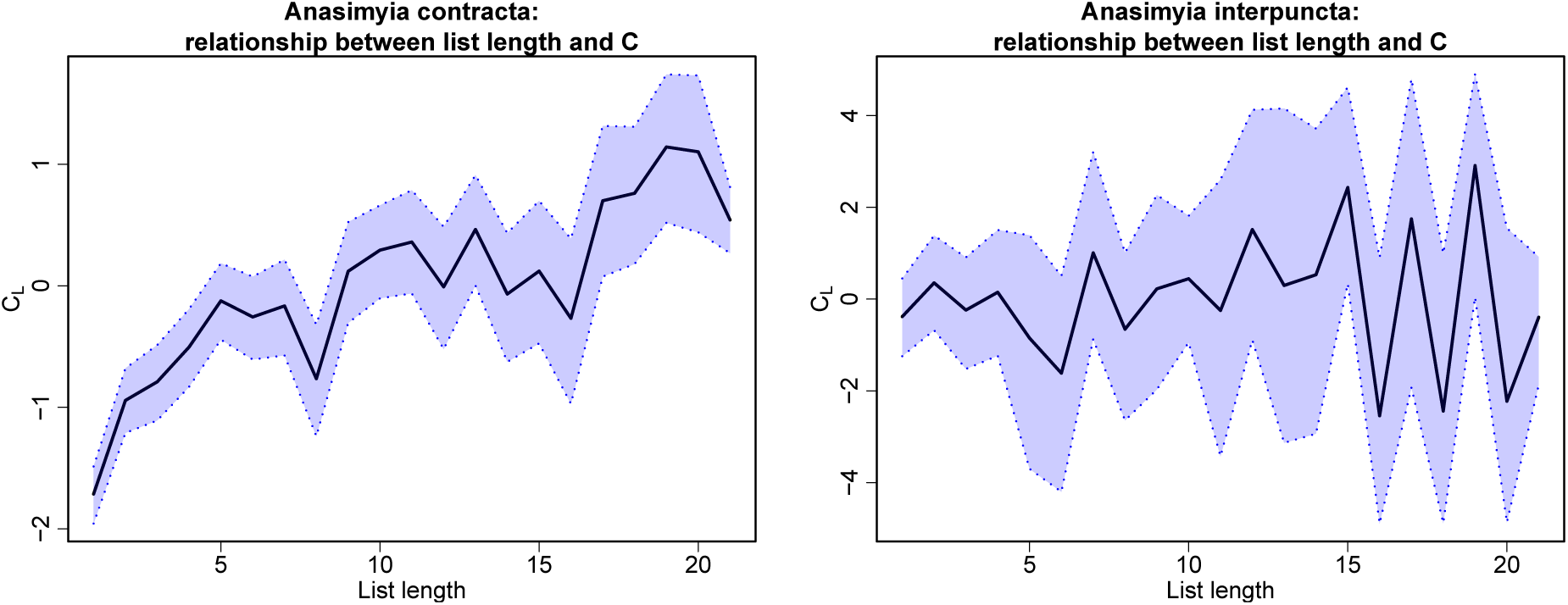
The solid black lines show the relationship between the list length and the categorical variable *C*_*L*_. The blue shaded areas show the 95% credible interval of the relationships. The left hand side plot uses the dataset of anasimyia contracta and the right hand site plot uses the dataset of anasimyia interpuncta.

From Figure 12, the plot of the relationship for anasimyia contracta is similar to the logarithm plot while the relationship between the list length and the *C*_*L*_ is obviously not logarithmic. Therefore, the relationship between the list length and the probability of detection varies between species. As mentioned in Section 1.2, the list length can represent the effort of searching the species. Then, these two plots show that the more efforts of searching the species does not mean the larger probability the species will be detected. For example, if there is a species afraid of people (e.g. yeti), no matter how many efforts used to search this species, the probability of detection will not increase.

### 5.3 Model complexity and fit

As mentioned in Section 4.3, DIC is a tool to compare the complexity and fit of models and the model is usually better if the value of DIC is lower. Hence, the table with DIC values of all models for anasimyia contracta and anasimyia interpuncta is shown in Table 3.

**Table 3:**
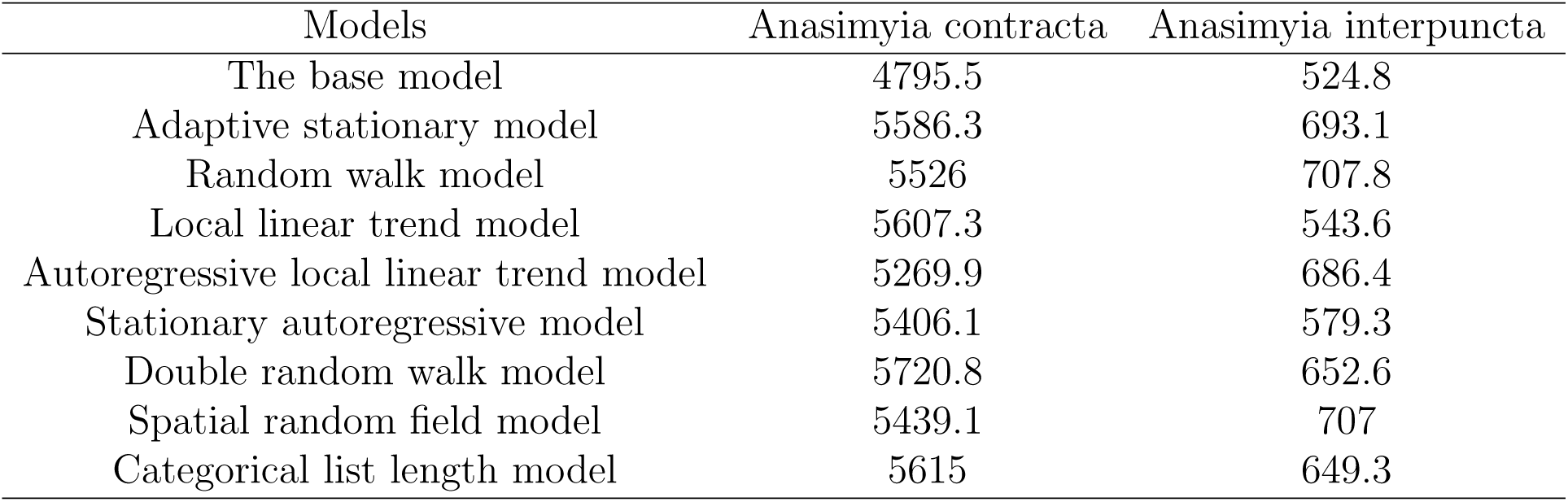
The DIC values of all models for anasimyia contracta and anasimyia interpuncta datasets.

| Models | Anasimyia contracta | Anasimyia interpuncta |
| --- | --- | --- |
| The base model | 4795.5 | 524.8 |
| Adaptive stationary model | 5586.3 | 693.1 |
| Random walk model | 5526 | 707.8 |
| Local linear trend model | 5607.3 | 543.6 |
| Autoregressive local linear trend model | 5269.9 | 686.4 |
| Stationary autoregressive model | 5406.1 | 579.3 |
| Double random walk model | 5720.8 | 652.6 |
| Spatial random field model | 5439.1 | 707 |
| Categorical list length model | 5615 | 649.3 |

If we only use the values of DIC to compare all models, the base model has the lowest DIC value for both species, which means that the base model has the best complexity and fit among all models, but from the plot for the base model in Section 5.2, we can see that the base model has the worst performance among all models.

In addition, from the values of DIC, the stationary autoregressive model performs well for both species while the random walk model, the adaptive stationary model and the double random walk model do not perform well for these species. However, from the plots in Section 5.2, the random walk model performs well when estimating the proportion of occupied sites for both species.

Furthermore, other models have different performances for different species if we compare the values of DIC. The autoregressive local linear trend model and the spatial random field model perform well for anasimyia contracta but not for anasimyia interpunta, whereas the local linear trend model and the categorical list length model perform well for anasimyia interpuncta, but not for anasimyia contracta.

However, the results from Section 5.2 prove that the DIC value is not suitable for comparing these models. Both plots and the simulation experiment from Outhwaite et al. (2018) show that the base model performs the worst and the random walk model performs the best among the first three models. However, the comparison of the values of DIC derives the opposite conclusions. Therefore, we can conclude that the comparison of DIC values is not suitable in this case.

### 5.4 Conclusions

In this chapter, we used three methods to compare the performance of each model. Firstly, from the convergence situation of each model, the local linear trend model, the autoregressive local linear trend model and the double random walk model are hard toconverge. Also, the spatial random field model used a long time to converge which is really time-consuming. Therefore, these models are not suitable to estimate the trend of the species in a short time.

Next, from the plot of each model, the trends of the species using the base model, the adaptive stationary model and the stationary autoregressive model fluctuated greatly from year to year. Also, the credible intervals of the base model are extremely wide. Then, from the trend estimated using each model, the adaptive stationary model, the stationary autoregressive model and the double random walk model tend to estimate the trend flatly, especially for the species anasimyia interpuncta. Besides, although introducing the slope *β*_*t*_ in the local linear trend model and the autoregressive local linear trend model is to show the trend directly from the slope, the results of the slopes in both models do not match the trends shown in their plots. This may be caused by their lack of convergence. In addition, the random walk model and the categorical list length model show similar trend for both species. However, the plot of the relationship between the list length and the variable *C*_*L*_ shows that the logarithmic relationship between these two quantities in the structures is not correct. Therefore, the categorical list length model will be more precise than the random walk model although their estimations are similar when fitting the anasimyia contracta and anasimyia interpuncta datasets.

Finally, from the comparison of the DIC values of all models, we can see that the results are totally different from the results evaluated from the plots. Therefore, the DIC value is not suitable for comparing the performance of each model.

As a result, the categorical list length model is the best model from the comparisons, because this model takes a reasonable time to converge and the result is the most precise among all models.

## 6 Application of the model to all sites

In previous chapters, we reviewed three occupancy models, suggested six potential improvements to the occupancy model and evaluated the performance of each model using the anasimyia contracta and anasimyia interpuncta datasets with only the informative sites. Then, in Section 5.4, we concluded that the categorical list length model had the best performance among all models. In this chapter, we will use the dataset of episyrphus balteatus and eristalis pertinax introduced in Section 1.2 to fit the categorical list length model including all sites. Because these are more realistic examples than those of the previous chapters. The reason of choosing these two particular species is that they are widespread species and therefore will have higher proportions of occupied sites, which will show the trend in the plots clearly. If we choose a scarce species, the proportions will be close to zero and hence the lower bound of the credible interval will tend to zero, which will be difficult to see the trend of the species clearly from the plot. Besides, we only use the data from the last 20 years to fit the model because the dataset is too large so that it will take several days to get the result for dataset with 45 years. The results of these species will be shown in this chapter.

### Episyrphus balteatus

The episyrphus balteatus dataset for the last 20 years is fitted using the categorical list length model. 10,000 MCMC iterations are run with the first 6,000 iterations discarded as burn-in. The running time is around 13 hours using JAGS through R.

The plot of the posterior mean of the proportion of occupied sites for episyrphus balteatus with 95% credible interval is shown on the left hand side of Figure 13. The proportion of occupied sites of the species varies frequently from year to year.

**Figure 13:**
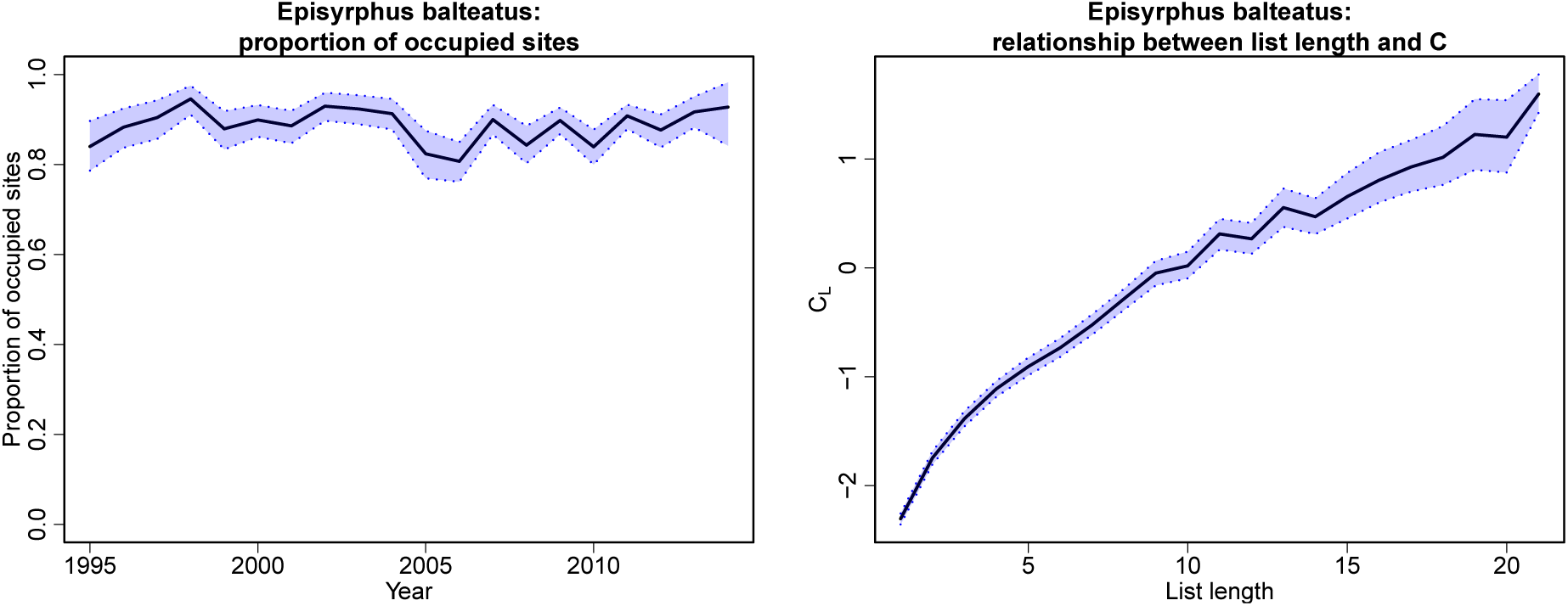
The plot on the left hand side shows the trend of the episyrphus balteatus estimated by the categorical list length model. The solid black line shows the posterior mean of the proportion of occupied sites from 1995 to 2014. The blue shaded area shows the 95% credible interval of the proportion. The plot on the right hand side shows the relationship between the list length and the categorical variable *C*_*L*_. The blue shaded area shows the 95% credible interval of the relationship.

To monitor the progress of the biodiversity, indicators are calculated both in the short term (5 years) and in the long term (20 years). The indicator of the episyrphus balteatusin the short term is calculated by the difference between the proportion of occupied sites in 2010 and 2014 which is used by Department for Environment, Food and Rural Affairs to calculate the indicators (Department for Environment, Food and Rural Affairs, 2017). Hence, the indicator of the episyrphus balteatus in the short term is 8.83%, which means the range of the species increased a lot from 2010 to 2014. However, there are fluctuations in the trend of the species, therefore, the indicator is sensitive to the choice of starting year. In order to reduce the effect of extreme years, the indicator in the long term is calculated by the difference between the mean of the proportion in the latest three years and that in the beginning three years. Hence, the indicator for the episyrphus balteatus in the long term is 3.13%, which means the range of the species also increased in the long term.

Then, we can use the categorical list length model to check the relationship between the list length and the detection probability. The plot of the relationship between these two quantities of the episyrphus balteatus with 95% credible interval is shown on the right hand side of Figure 13. Therefore, as more sampling efforts put into searching the species,the probability of detection of episyrphus balteatus increases.

### Eristalis pertinax

Similarly, the eristalis pertinax dataset for the last 20 years is fitted by the categorical list length model with 10,000 MCMC iterations and the first 6,000 iterations are discarded as burn-in. This process takes about 13 hours.

The plot of the posterior mean of the proportion of occupied sites for eristalis pertinax with 95% credible interval is shown on the left hand side of Figure 14. The overall proportion of occupied sites are very high, which is around 0.9. The proportions do not show much increase or decrease over the last 20 years, which means the range of the species remains stable.

**Figure 14:**
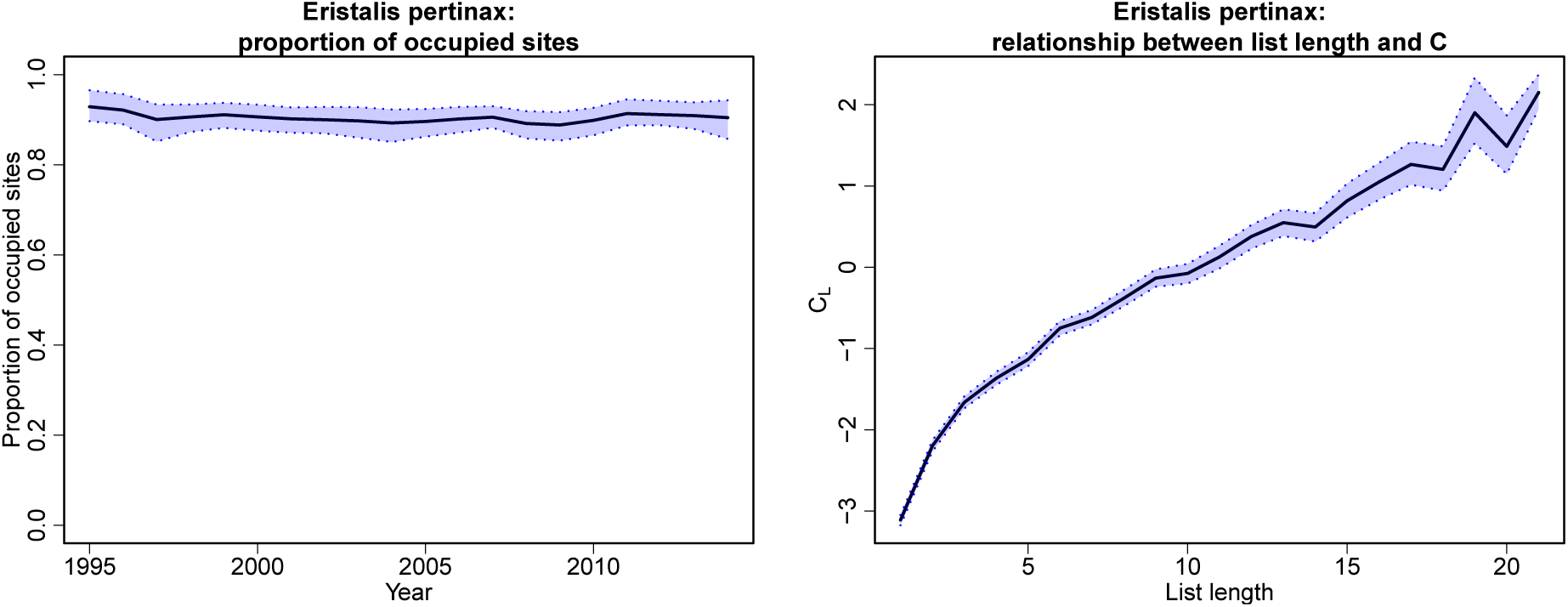
The plot on the left hand side shows the trend of the eristalis pertinax estimated by the categorical list length model. The solid black line shows the posterior mean of the proportion of occupied sites from 1995 to 2014. The blue shaded area shows the 95% credible interval of the proportion. The plot on the right hand side shows the relationship between the list length and the categorical variable *C*_*L*_. The blue shaded area shows the 95% credible interval of the relationship.

Next, the indicators of eristalis pertinax in the short term and in the long term are calculated using the same methods as before. Therefore, the the indicator for the species in the short term is 0.60% and that in the long term is 0.86%, so the range of the species remains stable with slight increases both in the short term and in the long term.

Finally, from the plot on the right hand side of Figure 14, there is a logarithmic relationship between the list length and the categorical variable *C*_*L*_. Therefore, as the sampling effort grows, the detection probability of eristalis pertinax also increases.

## 7 Discussion

This paper began by introducing the background related to monitoring the biodiversity. Then we described the datasets of the species were used to evaluate the models. From the dataset, if there is no record for a species in a particular site, we do not know whether it is because the species is not there, or because the species is present but not recorded. Therefore, a robust model is needed to estimate the trend of the species.

Hence, we reviewed three occupancy-detection models in Outhwaite et al. (2018). The base model has currently been used by ecologists to analyse the species, but from the results of this model for the species anasimyia contracta and anasimyia interpuncta, the base model was not suitable for estimating their trends because the trends estimated by these models fluctuated violently with wide credible intervals. Then the improvement of the base model was the adaptive stationary model. Although this model was much better than the previous model, the estimated trends still fluctuated greatly from year to year and the overall trends were flat which could not show the real trend of the species. The random walk model performed well across all criteria for both species. Therefore, the random walk model was recommended by Outhwaite et al. (2018) as being suitable for analysing the low recording intensity data.

However, there are still some opportunities to improve the random walk model further. Therefore, six alternative models were considered in this paper, with each alternative designed to improved on some aspects of the original random walk model. The local linear trend model and the autoregressive local linear trend model introduced the slope *β*_*t*_ to show the trend of the species directly. However, the MCMC were difficult to converge for these two models and therefore the slopes could not show the real trends of the species, particularly for the anasimyia interpuncta dataset which had fewer records. Then the stationary autoregressive model replaced the random walk process in the random walk model by the first order autoregressive process so that this model was stationary. However, the trends produced by this model fluctuated more violently than that by the random walk model. Then, the double random walk model applied the random walk process to both year effects, however, this model was hard to converge. Next, the spatial random field model considered the dependence between sites, but this model needed a long running time to fit the data due to using another software. Finally, the categorical list length model changed the structure of the random walk model by changing the relationship between the detection probability and the list length. The running time of this model was reasonable and the estimated trends were similar with those by the random walk model. Moreover, this model revealed the relationship between the list length and the detection probability.

However, when running a large dataset using these models, it takes a long time. Hence, massive computer resources are need to apply this model to a large dataset. Therefore, further works can be done by developing a simpler model which is more appropriate for routine application to large datasets.

Further improvement could also be gained by adding the seasonal effect or the month effect, since the species may have different probabilities to be detected in different seasons or months. For example, as introduced in Section 1.2, the flight period of the anasimyia contracta is in summer. Therefore, it would be easier to find this species in summer than in any other seasons. However, there will be additional computational burden because there are more parameters in the model.

It is also worth trying to change another way to record the location of detection of the species. Now, the location is recorded according to the site on a 1 *km*^2^ grid. This could be changed by recording based on the types of the lands, such as urban areas, forests, wetlands and so on. This is because the species may prefer staying in one or several particular types of lands. For example, in Section 1.2, the anasimyia contracta lives in the wetlands and ponds. Therefore, it is more likely to find this species near wetlands and ponds. Besides, it will also reduce the computational burden: in the spatial random field model, there are more than 9,000 big square sites, so it takes an extremely long running time to get a similar result with that of the random walk model. Since the number of the land types is much smaller than 9,000, the running time would be reduced.

## Acknowledgements

We would like to acknowledge the British Hoverfly Recording Scheme (BHRS) who provided the data used in this research. We are particularly grateful to Roger Morris of the BHRS for answering several queries about the data. Thanks are also due to Charlotte Outhwaite (UCL Department of Genetics, Evolution and Environment) for providing the organised data and helping with the queries about the data.

## Appendix

The appendix provides the code of the model definition of each model wrote in R.

~~~
Objects in these models:

DATA
====
nyear       Number of years in dataset
nsite       Number of sites in dataset
nvisit      Number of visits in dataset
y[j]        Detection status for the jth visit in the dataset
logL[j]     Logarithm of list length for jth visit in dataset
Site[j]     Site associated with the jth visit in the dataset
Year[j]     Year associated with the jth visit in the dataset

UNKNOWNS (i.e. quantities for which initial values are needed)
==========
z[i,t]       True occupancy status for site i in year t (binary).
b[t]         Year effect in occupancy model.
Beta[t]      Slope of the year effect in occupancy model.
phi          Parameter of the autoregressive process of the year effect.
u[i]         Site effect in occupancy model.
a[Year[j]]   Year effect in detection model for year associated with visit j.
c            Coefficient of logL in detection model, sampling effort effect.

OTHER QUANTITIES (internal use only)
================
psi[i,t]          Defined as P(z[i,t]=1).
Py[j]             Defined as P(y[j]=1).
p[j]              P(y[j]=1|z[i]=1).
mu.*              Mean in prior for parameter *.
tau.*             Precision in prior for parameter *.
sd.*              Prior standard deviation for parameter *.
if_branch[j] Binary variable, equal to 0 if list length greater than 20, equal to 1 otherwise.
L_small_10[j] Equal to list length if list length is not greater than 20, equal to 21 otherwise.

DERIVED PARAMETERS
==================
psi.fs              Proportion of occupied sites - model output
~~~

### The base model

~~~
function() {
   ### Priors ###
# State model priors
 for(t in 1:nyear){
  b[t] ˜ dunif(-10,10)     	# fixed year effect
}
for (i in 1:nsite) {
  u[i] ˜ dnorm(0, tau.u)   	# random site effect
}
tau.u <- 1/(sd.u * sd.u)
sd.u ˜ dt(0, 1, 1)%_%T(0,) 	#   half-Cauchy hyperpriors

	# Observation model priors
for (t in 1:nyear) {
  a[t] ˜ dnorm(mu.a, tau.a)   	# random year effect
}
mu.a ˜ dnorm(0, 0.01)
tau.a <- 1 / (sd.a * sd.a)
sd.a ˜ dt(0, 1, 1)%_%T(0,) 	#   half-Cauchy hyperpriors
c ˜ dunif(-10, 10) 	# sampling effort effect
##	# Model ###
	# State model
for (i in 1:nsite){
  for (t in 1:nyear){
   z[i,t] ˜ dbern(psi[i,t])
   logit(psi[i,t])<- b[t] + u[i]
}}
	# Observation model
for(j in 1:nvisit) {
  y[j] ˜ dbern(Py[j])
  Py[j]<- z[Site[j],Year[j]]*p[j]
  logit(p[j]) <- a[Year[j]] + c*logL[j]
}
##	# Derived parameters ###
	# Finite sample occupancy - proportion of occupied sites
for (t in 1:nyear) {
  psi.fs[t] <- sum(z[1:nsite,t])/nsite
  }
}
~~~

### Adaptive stationary model

~~~
function() {
  ##	# Priors ###
	# State model priors
 for(t in 1:nyear){
 b[t] ˜ dnorm(mu.b, tau.b) 	# random year effect
}
mu.b ˜ dnorm(0, 0.01)
tau.b <- 1/(sd.b * sd.b)
sd.b ˜ dt(0, 1, 1)%_%T(0,) 	# half-Cauchy hyperpriors

for (i in 1:nsite) {
u[i] ˜ dnorm(0, tau.u) 	# random site effect
}
tau.u <- 1/(sd.u * sd.u)
sd.u ˜ dt(0, 1, 1)%_%T(0,) 	# half-Cauchy hyperpriors

	# Observation model priors
for (t in 1:nyear) {
   a[t] ˜ dnorm(mu.a, tau.a) 	# random year effect
}
mu.a ˜ dnorm(0, 0.01)
tau.a <- 1 / (sd.a * sd.a)
sd.a ˜ dt(0, 1, 1)%_%T(0,) 	# half-Cauchy hyperpriors
c ˜ dunif(-10, 10) 	# sampling effort effect

##	# Model ###

	# State model
for (i in 1:nsite){
  for (t in 1:nyear){
   z[i,t] ˜ dbern(psi[i,t])
   logit(psi[i,t]) <- b[t] + u[i]
   }}

	# Observation model
for(j in 1:nvisit) {
  y[j] ˜ dbern(Py[j])
  Py[j]<- z[Site[j],Year[j]]*p[j]
  logit(p[j]) <- a[Year[j]] + c*logL[j]
}

##	# Derived parameters ###
	# Finite sample occupancy - proportion of occupied sites
for (t in 1:nyear) {
  psi.fs[t] <- sum(z[1:nsite,t])/nsite
 }
}
~~~

### Random walk model

~~~
function() {
##	# Priors ###

  	# State model priors
  b[1] ˜ dnorm(mu.b, 0.0001) 	# random walk prior on year effect
  for(t in 2:nyear){
  b[t] ˜ dnorm(b[t-1], tau.b)
  }
  mu.b ˜ dnorm(0, 0.01)
  tau.b <- 1/(sd.b * sd.b)
  sd.b ˜ dt(0, 1, 1)%_%T(0,) 	# half-Cauchy hyperpriors

  for (i in 1:nsite) {
    u[i] ˜ dnorm(0, tau.u) 	# random site effect
  }
  tau.u <- 1/(sd.u * sd.u)
  sd.u ˜ dt(0, 1, 1)%_%T(0,) 	# half-Cauchy hyperpriors

  	# Observation model priors
  for (t in 1:nyear) {
    a[t] ˜ dnorm(mu.a, tau.a) 	# random year effect
  }
  mu.a ˜ dnorm(0, 0.01)
  tau.a <- 1 / (sd.a * sd.a)
  sd.a ˜ dt(0, 1, 1)%_%T(0,) 	# half-Cauchy hyperpriors
  c ˜ dunif(-10, 10) 	# sampling effort effect

  ##	# Model ###
  	# State model
  for (i in 1:nsite){
  for (t in 1:nyear){
    z[i,t] ˜ dbern(psi[i,t])
    logit(psi[i,t])<- b[t] + u[i]
  }}
  	# Observation model
  for(j in 1:nvisit) {
    y[j] ˜ dbern(Py[j])
    Py[j]<- z[Site[j],Year[j]]*p[j]
    logit(p[j]) <- a[Year[j]] + c*logL[j]
  }
  ##	# Derived parameters ###
  	# Finite sample occupancy - proportion of occupied sites
  for (t in 1:nyear) {
    psi.fs[t] <- sum(z[1:nsite,t])/nsite
  }
}
~~~

### Local linear trend model

~~~
function() {
  ##	# Priors ###
  	# State model priors
  b[1] ˜ dnorm(mu.b[1], 0.0001) 	# local linear trend prior on year effect
  Beta[1] ˜ dnorm(mu.beta, 0.0001)
  for(t in 2:nyear){
    mu.b[t] <- b[t-1] + Beta[t]
    b[t] ˜ dnorm(mu.b[t], tau.b)
    Beta[t] ˜ dnorm(Beta[t-1], tau.beta)
  }
  mu.b[1] ˜ dnorm(0, 0.01)
  mu.beta ˜ dnorm(0, 0.01)
  tau.b <- 1/(sd.b * sd.b)
  tau.beta <- 1/(sd.beta * sd.beta)
  sd.b ˜ dt(0, 1, 1)%_%T(0,) 	# half-Cauchy hyperpriors sd.beta ˜ dt(0, 1, 1)%_%T(0,)
  for (i in 1:nsite) {
    u[i] ˜ dnorm(0, tau.u) 	# random site effect
  }
  tau.u <- 1/(sd.u * sd.u)
  sd.u ˜ dt(0, 1, 1)%_%T(0,) 	# half-Cauchy hyperpriors
  	# Observation model priors
  for (t in 1:nyear) {
    a[t] ˜ dnorm(mu.a, tau.a) 	# random year effect
  }
  mu.a ˜ dnorm(0, 0.01)
  tau.a <- 1 / (sd.a * sd.a)
  sd.a ˜ dt(0, 1, 1)%_%T(0,) 	# half-Cauchy hyperpriors
  c ˜ dunif(-10, 10) 	# sampling effort effect

  ##	# Model ###
  	# State model
  for (i in 1:nsite){ for (t in 1:nyear){
    z[i,t] ˜ dbern(psi[i,t])
    logit(psi[i,t])<- b[t] + u[i]
  }}
  	# Observation model
  for(j in 1:nvisit) {
    y[j] ˜ dbern(Py[j])
    Py[j]<- z[Site[j],Year[j]]*p[j]
    logit(p[j]) <- a[Year[j]] + c*logL[j]
  }
  ##	# Derived parameters ###
  	# Finite sample occupancy - proportion of occupied sites for (t in 1:nyear) {
    psi.fs[t] <- sum(z[1:nsite,t])/nsite
  }
}
~~~

### Autoregressive local linear trend model

~~~
function() {
  ##	# Priors ###

  	# State model priors
  b[1] ˜ dnorm(mu.b[1], 0.0001)
          	# local linear trend with autoregressive prior on year effect
  Beta[1] ˜ dnorm(mu.beta[1], 0.0001)
  phi ˜ dunif(-0.9999,0.9999)
  for(t in 2:nyear){
    mu.b[t] <- b[t-1] + Beta[t]
    b[t] ˜ dnorm(mu.b[t], tau.b)
    mu.beta[t] <- Beta[t-1] * phi
    Beta[t] ˜ dnorm(mu.beta[t], tau.beta)
  }
  mu.b[1] ˜ dnorm(0, 0.1)
  mu.beta[1] ˜ dnorm(0, 0.1)
  tau.b <- 1/(sd.b * sd.b)
  tau.beta <- 1/(sd.beta * sd.beta)
  sd.b ˜ dt(0, 1, 1)%_%T(0,) 	# half-Cauchy hyperpriors sd.beta ˜ dt(0, 1, 1)%_%T(0,)
  for (i in 1:nsite) {
    u[i] ˜ dnorm(0, tau.u) 	# random site effect
  }
  tau.u <- 1/(sd.u * sd.u)
  sd.u ˜ dt(0, 1, 1)%_%T(0,) 	# half-Cauchy hyperpriors

  	# Observation model priors
  for (t in 1:nyear) {
    a[t] ˜ dnorm(mu.a, tau.a) 	# random year effect
  }

  mu.a ˜ dnorm(0, 0.1)
  tau.a <- 1 / (sd.a * sd.a)
  sd.a ˜ dt(0, 1, 1)%_%T(0,) 	# half-Cauchy hyperpriors
  c ˜ dunif(-10, 10) 	# sampling effort effect
  ##	# Model ###
  	# State model
  for (i in 1:nsite){
    for (t in 1:nyear){
      z[i,t] ˜ dbern(psi[i,t])
      logit(psi[i,t])<- b[t] + u[i]
    }}
  	# Observation model
  for(j in 1:nvisit) {
    y[j] ˜ dbern(Py[j])
    Py[j]<- z[Site[j],Year[j]]*p[j]
    logit(p[j]) <- a[Year[j]] + c*logL[j]
  }
  ##	# Derived parameters ###
  	# Finite sample occupancy - proportion of occupied sites
  for (t in 1:nyear) {
     psi.fs[t] <- sum(z[1:nsite,t])/nsite
  }
}
~~~

### Stationary autoregressive model

~~~
function() {
  ##	# Priors ###
  	# State model priors
  b[1] ˜ dnorm(mu.b[1], 0.0001) 	# autoregressive prior on year effect phi ˜ dunif(-0.9999, 0.9999)
  for(t in 2:nyear){
    mu.b[t] <- phi * b[t-1]
    b[t] ˜ dnorm(mu.b[t], tau.b)
  }
  mu.b[1] ˜ dnorm(0, 0.01)
  tau.b <- 1/(sd.b * sd.b)
  sd.b ˜ dt(0, 1, 1)%_%T(0,) 	# half-Cauchy hyperpriors
  for (i in 1:nsite) {
    u[i] ˜ dnorm(0, tau.u) 	# random site effect
  }
  tau.u <- 1/(sd.u * sd.u)
  sd.u ˜ dt(0, 1, 1)%_%T(0,) 	# half-Cauchy hyperpriors
  	# Observation model priors
  for (t in 1:nyear) {
    a[t] ˜ dnorm(mu.a, tau.a) 	# random year effect
   }
  mu.a ˜ dnorm(0, 0.01)
  tau.a <- 1 / (sd.a * sd.a)
  sd.a ˜ dt(0, 1, 1)%_%T(0,) 	# half-Cauchy hyperpriors
  c ˜ dunif(-10, 10) 	# sampling effort effect

  ##	# Model ###

  	# State model
  for (i in 1:nsite){
  for (t in 1:nyear){
    z[i,t] ˜ dbern(psi[i,t])
    logit(psi[i,t])<- b[t] + u[i]
}}
	# Observation model
for(j in 1:nvisit) {
  y[j] ˜ dbern(Py[j])
  Py[j]<- z[Site[j],Year[j]]*p[j]
  logit(p[j]) <- a[Year[j]] + c*logL[j]
}
##	# Derived parameters ###
	# Finite sample occupancy - proportion of occupied sites
for (t in 1:nyear) {
    psi.fs[t] <- sum(z[1:nsite,t])/nsite
  }
}
~~~

### Double random walk model

~~~
function() {
  ##	# Priors ###
  	# State model priors
  b[1] ˜ dnorm(mu.b, 0.0001) 	# random walk prior on year effect
  for(t in 2:nyear){
    b[t] ˜ dnorm(b[t-1], tau.b)
  }
  mu.b ˜ dnorm(0, 0.1)
  tau.b <- 1/(sd.b * sd.b)
  sd.b ˜ dt(0, 1, 1)%_%T(0,) 	# half-Cauchy hyperpriors
  
  for (i in 1:nsite) {
    u[i] ˜ dnorm(0, tau.u) 	# random site effect
  }
  tau.u <- 1/(sd.u * sd.u)
  sd.u ˜ dt(0, 1, 1)%_%T(0,) 	# half-Cauchy hyperpriors
  
  	# Observation model priors
  a[1] ˜ dnorm(mu.a, 0.0001)
  for (t in 2:nyear) {
    a[t] ˜ dnorm(a[t-1], tau.a) 	# random walk prior on year effect
  }
  mu.a ˜ dnorm(0, 0.1)
  tau.a <- 1 / (sd.a * sd.a)
  sd.a ˜ dt(0, 1, 1)%_%T(0,) 	# half-Cauchy hyperpriors
  c ˜ dunif(-10, 10) 	# sampling effort effect
  
  ##	# Model ###
  
  	# State model
  for (i in 1:nsite){
  for (t in 1:nyear){
    z[i,t] ˜ dbern(psi[i,t])
    logit(psi[i,t])<- b[t] + u[i]
   }}
 
  	# Observation model
  for(j in 1:nvisit) {
    y[j] ˜ dbern(Py[j])
    Py[j]<- z[Site[j],Year[j]]*p[j]
    logit(p[j]) <- a[Year[j]] + c*logL[j]
  }
  
  ##	# Derived parameters ###
  	# Finite sample occupancy - proportion of occupied sites
  for (t in 1:nyear) {
    psi.fs[t] <- sum(z[1:nsite,t])/nsite
  }
}
~~~

### Spatial random field model

~~~
function() {
   ##	# Priors ###
   
   	# State model priors
   b[1] ˜ dnorm(mu.b, 0.0001) 	# random walk prior on year effect
   for(t in 2:nyear){
     b[t] ˜ dnorm(b[t-1], tau.b)
   }
   mu.b ˜ dnorm(0, 0.1)
   tau.b <- 1/(sd.b * sd.b)
   sd.b ˜ dunif(0, 5) 	# half-uniform hyperpriors

   u[1:9100] ˜ car.normal(adj[],weights[],num[],tau.u)

   	# Markov random field prior on site effect
   tau.u <- 1/(sd.u * sd.u)
   sd.u ˜ dunif(0, 5) 	# half-uniform hyperpriors
   	# Observation model priors

   for (t in 1:nyear) {
    a[t] ˜ dnorm(mu.a, tau.a) 	# random year effect
   }
   mu.a ˜ dnorm(0, 0.01)
   tau.a <- 1 / (sd.a * sd.a)
   sd.a ˜ dunif(0, 5) 	# half-uniform hyperpriors
   c ˜ dunif(-10, 10) 	# sampling effort effect
   
 ##	# Model ###
 	# State model
 for (i in 1:9100){
   for (t in 1:nyear){
   z[i,t] ˜ dbern(psi[i,t])
   logit(psi[i,t])<- b[t] + u[i]
  }}
  	# Observation model for(j in 1:nvisit) {
   y[j] ˜ dbern(Py[j])
   Py[j]<- z[Site[j],Year[j]]*p[j]
   logit(p[j]) <- a[Year[j]] + c*logL[j]
 }
 
  ##	# Derived parameters ###
  	# Finite sample occupancy - proportion of occupied sites for (t in 1:nyear) {
  psi.fs[t] <- sum(z[1:9100,t])/9100
 }
}
~~~

### Categorical list length model

~~~
function() {
  ##	# Priors ###
  
  	# State model priors
  b[1] ˜ dnorm(mu.b, 0.0001) 	# random walk prior on year effect
  for(t in 2:nyear){
   b[t] ˜ dnorm(b[t-1], tau.b)
  }
  mu.b ˜ dnorm(0, 0.01)
  tau.b <- 1/(sd.b * sd.b)
  sd.b ˜ dt(0, 1, 1)%_%T(0,) 	# half-Cauchy hyperpriors
  for (i in 1:nsite) {
    u[i] ˜ dnorm(0, tau.u) 	# random site effect
  }
  tau.u <- 1/(sd.u * sd.u)
  sd.u ˜ dt(0, 1, 1)%_%T(0,) 	# half-Cauchy hyperpriors

  	# Observation model priors
  for (t in 1:nyear) {
     a[t] ˜ dnorm(mu.a, tau.a) 	# random year effect
  }
  mu.a ˜ dnorm(0, 0.01)
  tau.a <- 1 / (sd.a * sd.a)
  sd.a ˜ dt(0, 1, 1)%_%T(0,) 	# half-Cauchy hyperpriors
  for (k in 1:20) {
    c[k] ˜ dunif(-5, 5) 	# categorical sampling effort effect
  }
  c[21] <- -sum(c[1:20])
  ##	# Model ###
  	# State model
  for (i in 1:nsite){
  for (t in 1:nyear){
    z[i,t] ˜ dbern(psi[i,t])
    logit(psi[i,t]) <- b[t] + u[i]
  }}
  	# Observation model
   for(j in 1:nvisit) {
  if_branch[j] <- 1 - step(L[j] - 21)
  L_small_10[j] <- if_branch[j] * L[j] + 21 * (1 - if_branch[j]) y[j] ˜ dbern(Py[j])
    Py[j] <- z[Site[j],Year[j]]*p[j]
    logit(p[j]) <- a[Year[j]] + c[L_small_10[j]]
  }
  ##	# Derived parameters ###
  	# Finite sample occupancy - proportion of occupied sites for (t in 1:nyear) {
    psi.fs[t] <- sum(z[1:nsite,t])/nsite
  }
}
~~~

## References

Agresti, A. (1990). Categorical data analysis. New York ; Chichester : Wiley.

Baldock, K. C. R., Goddard, M. A., Hicks, D. M., Kunin, W. E., Mitschunas, N., Osgathorpe, L. M., Potts, S. G., Robertson, K. M., Scott, A. V., Stone, G. N., Vaughan, I. P., and Memmott, J. (2015). Where is the UK’s pollinator biodiversity? The importance of urban areas for flower–visiting insects. Proc. R. Soc. B, 282.

Best, N., Richardson, S., and Thomson, A. (2005). A comparison of Bayesian spatial models for disease mapping. Statistical methods in medical research, 14:35–59.

Burns, F., August, T., Eaton, M., Noble, D., Powney, G., and Isaac, N. (2017). UK Biodiversity Indicators 2017. Indicator C4a. Status of UK priority species: relative abundance. Technical report, Joint Nature Conservation Committee.

Chandler, R. E. and Scott, E. M. (2011). Statistical methods for trend detection and analysis in the environmental sciences. John Wiley & Sons.

Convention on Biological Diversity (2010). Strategic plan for biodiversity 2011-2020 and the Aichi targets. Available from https://www.cbd.int/sp/default.shtml.

Department for Environment, Food and Rural Affairs (2017). UK biodiversity indicators 2017. Available from jncc.defra.gov.uk/ukbi.

Encyclopædia Britannica (2018). Hover fly. Online.

Farquharson, K. A., Hogg, C. J., and Grueber, C. E. (2018). A meta-analysis of birth-origin effects on reproduction in diverse captive environments. Nature Communications, 9.

Gelman, A., Carlin, J. B., Stern, H. S., Dunson, D. B., Vehtari, A., and Rubin, D. B. (2013). Bayesian data analysis. Boca Raton : CRC Press, third edition.

Isaac, N. J. B., van Strien, A. J., August, T. A., de Zeeuw, M. P., and Roy, D. B. (2014). Statistics for citizen science: extracting signals of change from noisy ecological data. Methods in Ecology and Evolution, 5:1052–1060.

Lunn, D., Jackson, C., Best, N., Thomas, A., and Spiegelhalter, D. (2013). The BUGS book: a practical introduction to Bayesian analysis. CRC Press.

Melinscaka, F. and Montesanoa, L. (2016). Beyond p-values in the evaluation of brain-computer interfaces:A Bayesian estimation approach. Journal of Neuroscience Methods, 270:30–45.

Outhwaite, C., Powney, G., August, T., and Isaac, N. (2017). UK Biodiversity Indicators 2017. Indicator C4b. Status of UK priority species: distribution. Technical report, Joint Nature Conservation Committee.

Outhwaite, C. L., Chandler, R. E., Powney, G. D., Collen, B., Gregory, R. D., and Isaac, N. J. (2018). Prior specification in Bayesian occupancy modelling improves analysis of species occurrence data. Ecological Indicators, 93:333–343.

Plummer, M. (2003). JAGS: A program for analysis of Bayesian graphical models using Gibbs sampling. In Proceedings of the 3rd International Workshop on Distributed Statistical Computing, Vienna, Austria. pp. 20–2.

R Core Team (2015). R: A language and environment for statistical computing. R Foundation for Statistical Computing, Vienna, Austria.

Speight, M. C. D. (2011). Species accounts of European Syrphidae (Diptera), Glasgow 2011. In Syrph the Net, the database of European Syrphidae, volume 65. Syrph the Net publications, Dublin.

Spiegelhalter, D., Thomas, A., Best, N., and Lunn, D. (2014). OpenBUGS User Manual, Version 3.2.3. Online user manual, http://www.mrc-bsu.cam.ac.uk/bugs.

Spiegelhalter, D. J., Best, N. G., Carlin, B. P., and Van Der Linde, A. (2002). Bayesian measures of model complexity and fit. Journal of the Royal Statistical Society: Series B (Statistical Methodology), 64:583–6

Sturtz, S., Ligges, U., and Gelman, A. (2005). R2WinBUGS: A package for running WinBUGS from R. Journal of Statistical Software, 12(3):1–16.

Su, Y.-S. and Yajima, M. (2015). R2jags: Using R to run ‘JAGS’. R package version 0.5–7.

Szabo, J. K., Vesk, P. A., Baxter, P. W. J., and Possingham, H. P. (2010). Regional avian species declines estimated from volunteer-collected long-term data using List Length Analysis. Ecological Applications, 20(8):2157–2169.

